# The early-life determinant LIN28B constrains immunoglobulin light chain secondary rearrangements independent of BCR specificity

**DOI:** 10.64898/2026.09.04.749379

**Authors:** Elena Boldrin, Hugo Åkerstrand, Giorgia Montano, Niklas Segrén, Christine Valfridsson, Stefan Lang, Joan Escrivá-Font, Camila Rosat Consiglio, Shamit Soneji, Joan Yuan

**Affiliations:** Developmental Immunology Unit, Division of Molecular Hematology, Department of Laboratory Medicine, Lund Stem Cell Center, Lund University, Lund 22242, Sweden; Computational Genomics Unit, Division of Molecular Hematology, Department of Laboratory Medicine, Lund Stem Cell Center, Lund University, Lund 22242, Sweden; Systems Immunology Unit, Division of Molecular Hematology, Department of Laboratory Medicine, Lund Stem Cell Center, Lund University, Lund 22242, Sweden

## Abstract

The early-life B cell repertoire is disproportionately enriched for self-reactive specificities in mice and humans, raising the question of how this ontogenic permissiveness is achieved. The predominant B cell central tolerance mechanism edits away self-reactivity by secondary rearrangements of the immunoglobulin light chain (IgL) following strong B cell receptor (BCR) engagement during the immature B cell stage. Here, we demonstrate a layer of developmental regulation, imposed by the early-life restricted RNA-binding protein LIN28B, that suppresses the incidence and capacity for IgL secondary rearrangements during ontogeny. Genetic dissection demonstrated that the underlying mechanisms operate independent of BCR specificity or pre-BCR requirement, dissociating the receptor editing fate from strict BCR instruction. We identified an adult-specific receptor editing-biased pre-B cell state marked by CD25 expression and metabolic quiescence. LIN28B subverted this state, shifting the balance from secondary rearrangements to positive selection and bone marrow egress. Together, our results demonstrate that the central tolerance threshold is an ontogenically tuned parameter, providing insights into the self-reactivity bias that characterizes the early-life B cell repertoire.

**One Sentence Summary:** The developmentally restricted RNA-binding protein LIN28B limits the extent of Immunoglobulin light chain receptor editing to shift the balance from stringent self-tolerance towards accelerated B cell output early in life.

## INTRODUCTION

Fetal and neonatal B cell repertoires are known to be enriched for self-reactive and poly-reactive specificities with restricted diversity, in contrast to the more tightly censored repertoires of the adult(*1–3*). In mice, early-life B cell development is marked by an increased permissiveness toward the selection of self-reactive B-1 cell clones which do not efficiently emerge in adults(*4*). In humans, a high prevalence of CD5⁺ B cells in fetal and cord blood is associated with a high frequency of reactivity against self as well as microbial antigens(*5–7*). Together, these data are consistent with the notion that early-life B cell selection may have been shaped to retain certain evolutionarily useful self-reactive specificities(*8*), raising the question of how this permissiveness toward self-reactivity is achieved.

The central tolerance checkpoint serves as the principal barrier against self-reactivity. This checkpoint is enforced as newly formed immature B (immB) cells express surface IgM, whereby strong B cell receptor (BCR) engagement triggers secondary rearrangements of the immunoglobulin light chain (IgL), a process known as receptor editing, with clonal deletion being a less frequent fate(*9–13*). During the editing process immB cells re-express RAG1/2 in a reverse developmental trajectory, enabling pre-B cell stage-like rearrangements to purge self-reactivity, generally starting at the *Igκ* locus, with *Igλ* rearrangements engaged later (*14, 15*). Disruptions in receptor editing have been linked to various autoimmune diseases, such as systemic lupus erythematosus, type 1 diabetes, and myasthenia gravis(*16, 17*), emphasizing its essential role in enforcing self-tolerance. Additional checkpoints contribute to restrict self-reactivity both upstream, where poor heavy chain pairing with the surrogate light chain (SLC) limits the expansion of certain self-reactive clones at the pre-BCR stage(*18, 19*), and downstream, where residual self-reactive B cells that escape central tolerance are controlled by anergy in the periphery(*12, 20, 21*).

In the life of a B cell, differentiation programs are broadly organized to insulate RAG1/2-mediated V(D)J recombination from proliferative cellular states to protect cells from genotoxic stress and the risk of neoplastic transformation (*22–24*). Consistent with this principle, elevated PI3K/AKT/mTOR signaling triggered by either the BCR or CD19 drives an anabolic program that opposes the quiescent, recombination-permissive state (*25, 26*). Enhanced signaling through this pathway relaxes the central tolerance checkpoint and promotes the forward developmental trajectory of immB cells by simultaneously restricting secondary rearrangements and driving bone marrow egress. The former is achieved through the phosphorylation and degradation of FOXO1 required for *Rag1/2* expression(*27–29*), and the latter through downregulation of CXCR4, critical for pre-B cell identity, niche retention, and IgL locus accessibility (*30, 31*).

These mechanisms, however, have been characterized almost exclusively in adult bone marrow (BM), leaving it unclear how altered tolerance regulation in early life might permit the output of self-reactive B cells. One potential mechanism is the pre-BCR-independent generation of immB cells, where premature *Igk* recombination has been reported to allow for the developmental progression of SLC-incompatible self-reactive heavy chains that would otherwise be counter selected at the pre-BCR checkpoint(*18, 32*). While receptor editing is known to operate during neonatal life in both mice and humans (*7, 33*) how its regulation changes during ontogeny and whether early-life editing permissiveness could contribute to license self-reactive B cell output remain poorly understood.

A unique opportunity to interrogate the basis of altered tolerance regulation during early ontogeny is offered by the RNA-binding protein LIN28B, a key regulator of fetal and neonatal lymphopoiesis. LIN28B expression in hematopoiesis is restricted to fetal and neonatal life, downregulated by 3 weeks postnatally, and is both necessary and sufficient to drive hallmark features of neonatal B cell development(*34, 35*). These include enhanced positive selection of self-reactive B cells, including CD5⁺ B-1a cells(*36–38*). Mechanistically, LIN28B suppresses let-7 microRNA biogenesis, thereby de-repressing the downstream MYC-dependent transcription and augmenting PI3K-mediated tonic signaling(*37, 39*). The combined effect elevates overall protein synthesis and anabolic state during the pre-B and immB developmental stages(*35*). These findings establish LIN28B as a programmable molecular axis that imposes neonatal-like properties on B cell development.

Here, we exploit genetic models of LIN28B perturbation to interrogate how tolerance stringency is regulated during early ontogeny. We show that LIN28B-induced programming operates independently of the pre-BCR checkpoint and instead acts at the immB stage to limit IgL receptor editing. Remarkably, this effect is independent of BCR-specificity and accompanied by accelerated developmental progression and bone marrow egress. Our findings reveal that LIN28B uncouples positive selection from stringent BCR quality control by interfering with the metabolically quiescent state required for efficient receptor editing at the central tolerance checkpoint. We conclude that the rules of central tolerance are not fixed but rather ontogenically tuned, with consequences for the establishment of the neonatal peripheral B cell pool.

## RESULTS

### LIN28B reduces Immunoglobulin light chain recombining pre-B cells independently of the pre-BCR

LIN28B expression reshapes BM B cell development in early life, licensing the emergence of immB cells bearing markers of self-reactivity(*37, 38*). To dissect how LIN28B shapes B cell developmental progression we performed single-cell RNA sequencing (scRNAseq). CD19^+^B220^+^CD93^+^ B cell progenitors (BCPs) were isolated from the BM of doxycycline-treated control (ctrl, *Rosa26^rtTA*m2^*) and tet-LIN28B (*Col1a^tetO-LIN28B^ Rosa26^rtTA*m2^*) adult mice (n=2 per group), where ectopic LIN28B expression was induced with a doxycycline diet for 14 days. Recirculating B cells (CD19^+^B220^+^CD93^−^) were included at a frequency of 10% of input cells to anchor the developmental trajectory (Figure 1A). 18 200 high-quality cells were recovered across all samples. Uniform Manifold Approximation and Projection (UMAP) representation resolved the expected BCP populations, comprising cycling (phase S/G2/M) and recombining (*Rag1*⁺) pro-B cell clusters (clusters 1-2), cycling (phase S/G2/M) and recombining (*Rag1*⁺) pre-B clusters (*Igkc*⁺ or *Iglc*⁺; clusters 3-7), immB and recirculating mature B cell clusters (clusters 8-9) (Figure 1B-C, Supplementary Figure 1A-B). Interestingly, tet-LIN28B mice showed an increased representation of recombining pro-B cells, while the recombining pre-B cluster was reduced (cluster 7, pre-B_REC_) as compared to controls (Figure 1D-E, Supplementary Figure 1C-D). This difference was further reflected in the pseudotime trajectory analysis of BCPs from pre-B to immB cells, where control mice displayed a preferential accumulation at the pre-B_REC_ stage (Supplementary Figure 1E-F), consistent with LIN28B constraining the developmental window during which IgL recombination takes place. Flow cytometric distinction based on forward scatter into large (cycling) and small (recombining) pre-B populations failed to recapitulate the observed LIN28B induced reduction of the pre-B_REC_ subset frequency, highlighting the superior subsetting resolution provided by scRNAseq analysis (Supplementary Figure 1G).

**Figure 1.**
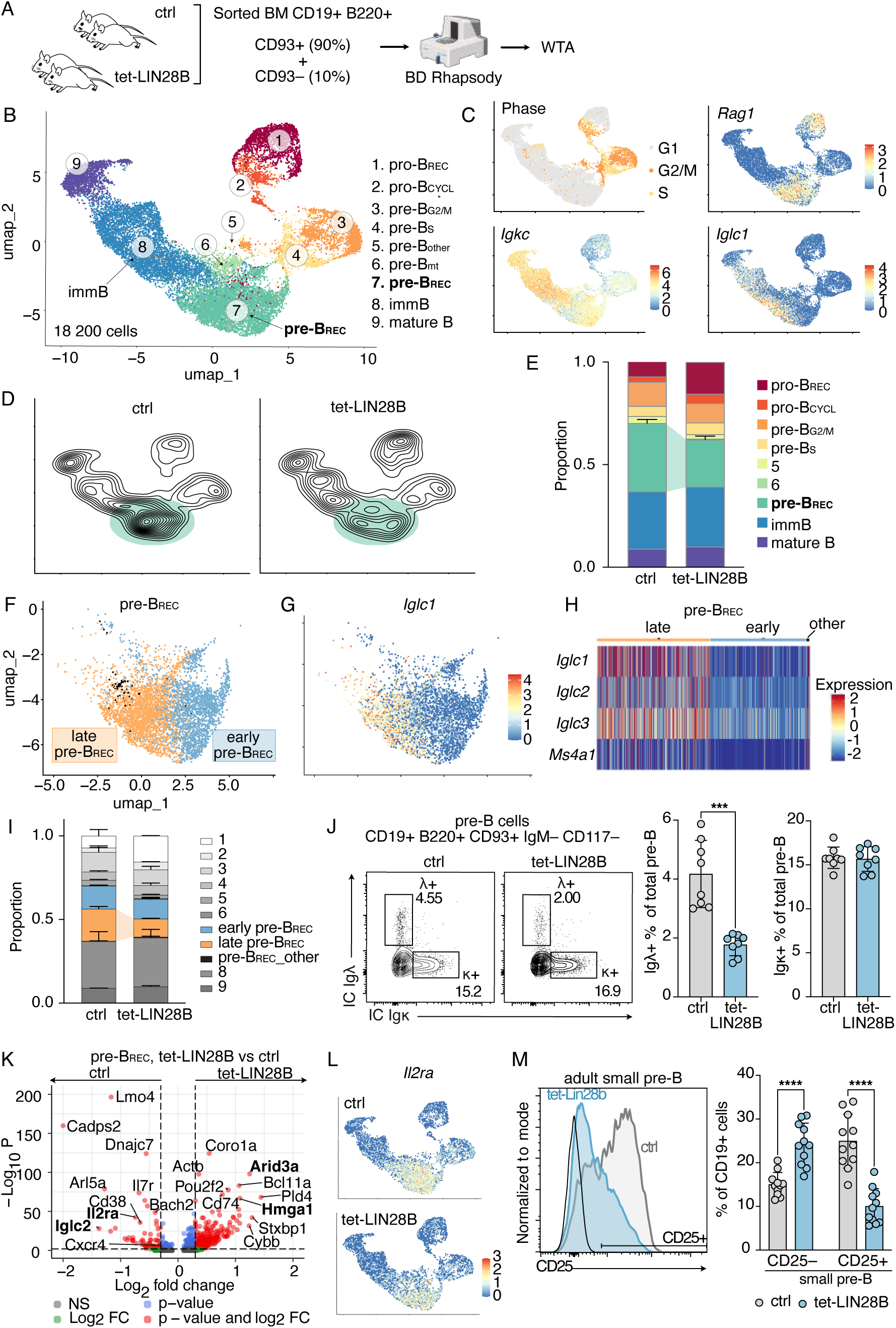
LIN28B reduces late recombining pre-B cells while limiting CD25 expression. A. Experimental layout of single-cell RNA sequencing (scRNAseq) of adult control (ctrl, n=2) and tet-LIN28B (n=2) mice. CD19⁺B220⁺CD93⁺ bone marrow (BM) cells were FACS sorted, mixed with 10% CD19⁺B220⁺CD93^−^ cells, and underwent scRNAseq by BD Rhapsody, followed by whole transcriptome analysis (WTA). B. Uniform Manifold Approximation and Projection (UMAP) representation of scRNAseq data with identified Seurat clusters: recombining pro-B (pro-B_REC_), cycling pro-B (pro-B_CYCL_), pre-B cells in G2/M (pre-B_G2/M_) or S phase (pre-B_S_), recombining pre-B (pre-B_REC_), immature B (immB) and mature recirculating B cells. C. Expression of selected B cell development genes and cell cycle phases according to scRNAseq data. D. Geometric density of ctrl and tet-LIN28B cells. The area including pre-B_REC_ cells is highlighted in green. E. Proportion of Seurat clusters in the different genotypes. Mean ± Standard Deviation (SD) shown for cluster 7/pre-B_REC_. F. UMAP of pre-B_REC_ cells, with Seurat clusters calculated upon reclustering (blue=early pre-B_REC_, orange=late pre-B_REC_, black=other pre-B_REC_). G. *Iglc1* expression in pre-B_REC_ cells. H. Heatmap of selected genes differentially expressed between early and late pre-B_REC_ cells. I. Proportion of early, late and other pre-B_REC_ Seurat clusters as identified in F in ctrl (n=2) and tet-LIN28B (n=2), shown as a fraction out of quality filtered total cells for each sample. Mean ±SD. J. Representative FACS plot and percentages of Igλ⁺ and Igκ⁺ out of pre-B cells (CD19⁺B220⁺CD93⁺IgM^−^CD117^−^) in ctrl (n=8) and tet-LIN28B (n=8) mice, identified by intracellular staining. Mean ±SD. Mann-Whitney test, *** p < 0.001, only significant comparisons shown. K. Volcano plot of differentially expressed genes in ctrl and tet-LIN28B pre-B_REC_ cells in the scRNAseq dataset. NS=non-significant, FC=fold change. L. *Il2ra* expression in ctrl and tet-LIN28B cells from the scRNAseq experiment. M. Representative FACS histogram of CD25 surface expression on small pre-B cells (CD19⁺B220⁺CD93⁺IgM^−^CD117^−^ FSC_low_) and percentages of CD25⁺ and CD25^−^ small pre-B cells out of CD19⁺ cells in ctrl (n=11) and tet-LIN28B (n=11) mice. Black line represents the fluorescence-minus-one control (FMO). Mean ±SD. Two-way ANOVA test, with Šídák correction for multiple comparisons, **** adjusted p-value (adj.p) < 0.0001.

Fetal B cell progenitors have been reported to be able to undergo premature *Igκ* chain recombination in the pro-B cell stage, bypassing the pre-BCR checkpoint and pre-B cell stages(*18, 32*). In line with this possibility, expression of *Igκ* germline (κ0) and recombined transcripts in pro-B cells from ctrl and tet-LIN28B mice was higher upon LIN28B expression consistent with enhanced locus opening and recombination (Supplementary Figure 2A-B). To directly test whether LIN28B acts through such a pre-BCR bypass mechanism, we next crossed the tet-LIN28B mice onto a surrogate light chain knockout background (SLC–/–)(*40*), deficient for *Vpreb1*, *Vpreb2* and *Igll1* and therefore unable to express a functional pre-BCR. LIN28B induction, however, failed to alleviate the SLC–/– imposed block in B cell development (Supplementary Figure 2C-D), arguing against pre-BCR bypass being a major mechanism of LIN28B action.

### LIN28B expression reduces late pre-B_REC_ cells while limiting CD25 expression

In the pre-B_REC_ stage, *Igκ* and *Igλ* recombination takes place during distinct and consecutive stages(*14, 15*). We therefore examined this population with higher resolution. Re-clustering yielded two major subsets with control cells enriched in a population characterized by the expression of *Igλ* locus transcripts (*Iglc1*, *Iglc2*, *Iglc3*, Figure 1F-H, Supplementary Figure 3A). Since the *Igλ* locus opens later than *Igκ* (*14*), often after *Igκ* has already undergone recombination attempts, we identified this population as late pre-B_REC_. Tet-LIN28B mice showed a marked and selective reduction of late pre-B_REC_ (Figure 1I, Supplementary Figure 3B-C), consistent with a reduction of cells undergoing secondary rearrangements. In line with this, intracellular Igλ protein expression was reduced in tet-LIN28B pre-B cells while intracellular Igκ expression remained comparable to controls (Figure 1J).

We next examined LIN28B-induced transcriptional signatures in the pre-B_REC_ cluster. Functional enrichment analysis revealed a modest increase in the expression of downstream targets of let-7, MYC and mTOR signaling (Figure 1K, Supplementary Table 1, Supplementary Figure 3D-E) consistent with its known roles(*36, 37, 41–44*). Interestingly, *Il2ra*/CD25 transcript levels were downregulated in tet-LIN28B pre-B_REC_ (Figure 1K-L, Supplementary Figure 3F). CD25 is a widely used surface marker of the pre-B cell stage, particularly highly expressed in IgL recombining pre-B cells(*45, 46*), though a functional role in this context has not been established. Consistent with this transcriptional change, FACS analysis confirmed reduced CD25 surface expression in the small pre-B cell stage of tet-LIN28B mice (Figure 1M). Thus, reduced late pre-B_REC_ representation and CD25 surface expression are hallmarks of LIN28B-induced B cell development.

### Neonatal B cell development exhibits reduced late pre-B_REC_ cells and CD25 expression

To investigate if the observed phenotype also occurred under physiological LIN28B expression in neonates, a separate scRNAseq of neonatal, adult ctrl and adult tet-LIN28B CD19⁺B220⁺CD93⁺ BCPs was performed using a targeted 463-transcript panel (*47*) (Figure 2A, Supplementary Table 2). Following cluster generation and identification, we found that neonatal BCPs showed a dramatic reduction of *Igλ* transcript positive late pre-B_REC_ cells (Figure 2B-E, Supplementary Figure 3G) compared to adult control BCPs. Tet-LIN28B adult BCPs exhibited an intermediate phenotype, indicating that LIN28B only partially recapitulates the neonatal phenotype and that other intrinsic and environmental factors are likely involved in ontogenic differences in B cell development. Similar to the adult, the observed changes in pre-B_REC_ representation by LIN28B were mild when assessed for small pre-B cell frequency by flow cytometry (Supplementary Figure 3H). scRNAseq and FACS analysis did however confirm a similar *Lin28b* dose-dependent *Il2ra*/CD25 suppression in neonatal pre-B cells (Figure 2F-G). Taken together, our findings demonstrate that neonatal B cell development is characterized by a reduced representation of late pre-B_REC_ cells and reduced CD25 expression, both of which are recapitulated by ectopic LIN28B expression.

**Figure 2.**
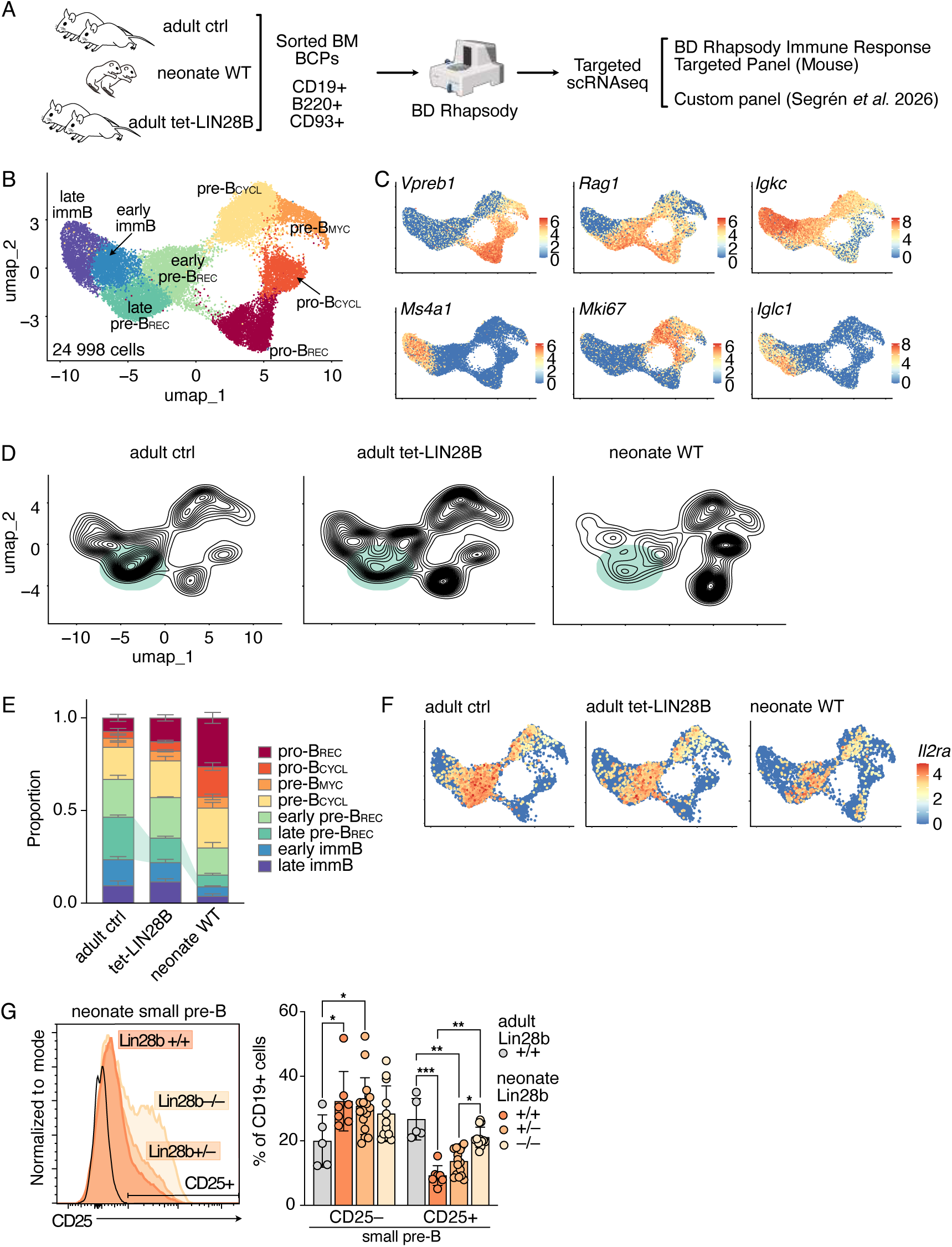
Neonatal B cell development exhibits reduced late pre-B_REC_ cells and CD25 expression. A. Experimental design of scRNAseq analysis of BCPs from adult ctrl (n=2), tet-LIN28B (n=2) and wildtype (WT) neonate (n=2 litters) mice: BM CD19⁺B220⁺CD93⁺ cells were FACS sorted and subjected to targeted scRNAseq using a targeted panel of genes including the BD Rhapsody Immune Response Targeted Panel (Mouse) and a custom panel(*47*). After filtering, 24998 high-quality cells were obtained and used for downstream analyses. B. UMAP representation of scRNAseq data with identified Seurat clusters. Clusters were defined as follows: pro-B cells (*Vpreb1*⁺) were separated into pro-B_REC_ (*Rag1*⁺) and pro-B_CYCL_ (*Mki67*⁺). Pre-B cells (*Vpreb1*^−^) were separated into pre-B_CYCL_ (*Mki67*⁺), pre-B_MYC_ (*Myc*), and pre-B_REC_ (*Rag1*⁺). ImmB cell clusters expressed *Ms4a1*. C. Expression of selected B cell development genes in scRNAseq data. D. Geometric density of adult ctrl and tet-LIN28B, and neonate WT cells. E. Proportion of Seurat clusters in ctrl adult (n=2), tet-LIN28B adult (n=2), and WT neonate (n=2) mice shown as fraction out of total quality filtered cells for each sample. Mean ±SD. F. *Il2ra* expression in adult ctrl, tet-LIN28B and WT neonate cells from the scRNAseq experiment. G. Representative FACS histogram of CD25 surface expression on small pre-B cells and percentages of CD25⁺ and CD25^−^ small pre-B cells out of CD19⁺ cells in adult control (n=5), and neonate *Lin28b +/+* (n=7), +/– (n=16) or –/– (n=12) mice (postnatal day 2-3). Black line represents the FMO. Mean ±SD. Two-way ANOVA test, with Tukey correction for multiple comparisons, only significant comparisons shown, * adj.p < 0.05, ** adj.p < 0.01, *** adj.p < 0.001.

### LIN28B expression limits the extent of IgL secondary rearrangements and receptor editing

To investigate the basis for reduced *Igλ* transcript positive late pre-B_REC_ cells, we first assessed whether it reflected reduced locus accessibility upon LIN28B induction(*48*). We measured *Ig*λ*1* germline transcript (GLT) expression from FACS sorted BCPs by RT-qPCR. LIN28B induction did not decrease *Igλ1* GLT in large or small pre-B cells (Figure 3A), arguing against impaired locus accessibility. To examine whether LIN28B reduces the occurrence of secondary rearrangements, we turned our attention to the *Igκ* gene segment usage among immB cells. Developing B cells undergoing IgL recombination show a bias towards the sequential sampling of first more proximal *Jκ* segments, followed by distal segments, and finally *Igλ* segments(*49*). PCR analysis of *Jκ* segment usage revealed a trend towards a reduced *Jκ5*/*Jκ1* ratio in tet-LIN28B immB cells (Figure 3B-C), consistent with a preference for more proximal segments among newly selected IgM⁺ B cells (*50*). Concomitantly, LIN28B expression reduced Igλ positive immB cells (Figure 3D), collectively indicating that LIN28B expressing cells carry a reduced history of secondary rearrangements. To establish whether this phenotype represented a physiological feature of early-life BCR repertoire selection, we compared Igλ usage in immB cells from neonatal and adult mice by flow cytometry. Neonatal immB cells showed reduced Igλ usage relative to adults in a LIN28B dose-dependent manner (Figure 3E). We conclude that suppression of secondary rearrangements is a LIN28B instructed feature of neonatal B cell maturation.

**Figure 3.**
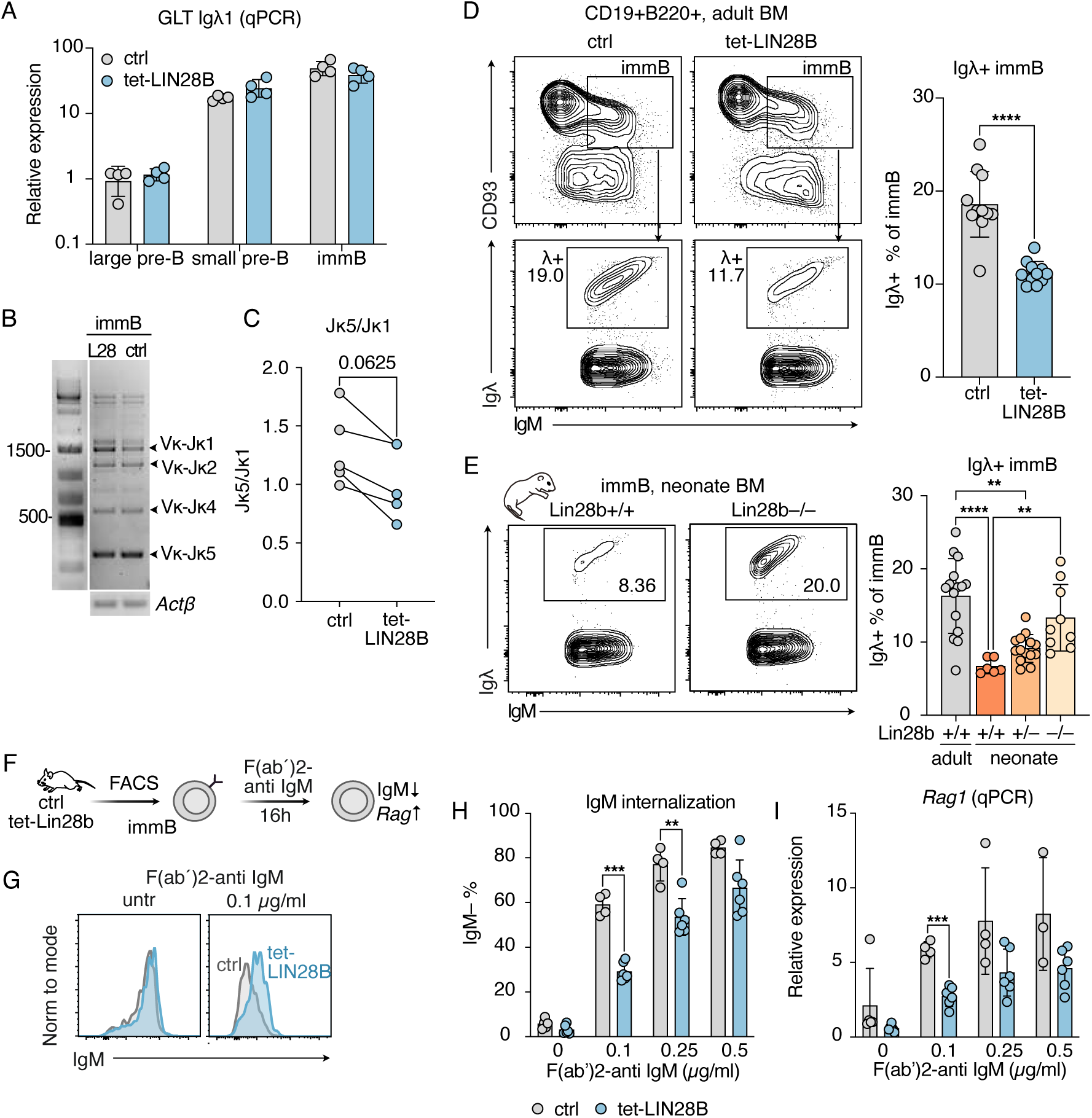
LIN28B expression limits the extent of IgL receptor editing early in life. A. *Igλ1* germline transcript (GLT) expression in sorted BCP populations from ctrl (n=4) and tet-LIN28B (n=4) assessed by qRT-PCR. Large pre-B: CD19⁺B220⁺CD93⁺IgM^−^CD117^−^FSC_high_; small pre-B: CD19⁺B220⁺CD93⁺IgM^−^CD117^−^ FSC_low_; immB: CD19⁺ B220⁺ CD93⁺ IgM⁺. Mean ±SD. Two-way ANOVA test, with Šídák correction. B. Representative agarose gel of PCR detecting *VκJκ* joints in sorted immB cells. C. Ratio of *Jκ5* over *Jκ1* usage in immB cells from quantification of PCR bands on agarose gel. Connecting lines link littermate ctrl and tet-LIN28B samples that were run in parallel on the same gel. Two-tailed Wilcoxon matched-pairs signed rank test. D. Representative FACS plot and percentages of Igλ⁺ out of immB cells in ctrl (n=11) and tet-LIN28B (n=11) mice, identified by surface staining. Mean ±SD. Mann-Whitney test. E. Representative FACS plot and percentages of Igλ⁺ out of immB cells in adult ctrl (n=15), and Lin28b +/+ (n=6), +/– (n=14) or –/– (n=9) neonate mice (postnatal day 2-3), identified by surface staining. Kruskal-Wallis test, with Dunn’s correction. F. Experimental layout of receptor editing induction: immB cells were FACS sorted and cultured *in vitro* in presence of increasing concentrations of F(ab′)2-anti IgM. Subsequently, surface IgM downregulation was measured by flow cytometry and *Rag1* induction by RT-qPCR. G. Representative FACS histogram of surface IgM expression of immB cells sorted from ctrl or tet-LIN28B mice after overnight culture with F(ab′)2-anti IgM (0.1 µg/ml) or left untreated (untr). H. Percentage of IgM^−^ cells in overnight culture of immB cells from ctrl (n=4) or tet-LIN28B (n=6) mice treated with the indicated concentrations of F(ab′)2-anti IgM. Mean ±SD. Two-way ANOVA test, with Šídák correction for multiple comparisons. I. *Rag1* expression assessed by qRT-PCR in cells in H. Expression was normalized to *Actb* and presented as relative to the average of the untreated ctrl samples from the same experiment. Mean ±SD. Two-way ANOVA test, with Šídák correction for multiple comparisons. Only significant comparisons shown, p/adj.p: * < 0.05, ** < 0.01, *** < 0.001, **** < 0.0001.

Secondary IgL rearrangements can occur following either non-productive recombination events or back-differentiation of immB cells triggered by self-antigen engagement during immB cell selection (*51, 52*). The latter is a process known as receptor editing and constitutes the main mechanism of central tolerance. To test whether the LIN28B-driven program dampens the receptor editing response, we FACS sorted tet-LIN28B and ctrl immB cells to induce receptor editing *ex vivo.* Cells were treated with F(ab’)2-anti IgM antibody, to model self-antigen mediated IgM crosslinking. After 16 hours, markers of receptor editing were induced in control cells in an anti-IgM dose dependent manner including downregulation of surface IgM and *Rag1* transcript upregulation (Figure 3F-I)(*51*). Interestingly, these hallmarks were diminished in tet-LIN28B immB cells relative to controls despite the lack of significant changes in cell viability (Supplementary Figure 4A). Taken together, our findings show that LIN28B limits the capacity for immB cells to undergo receptor editing.

### LIN28B suppresses secondary rearrangements of immB cells independently of IgH specificity

To address whether the LIN28B-programmed reduction in secondary rearrangements resulted from a skewed Ig heavy chain (IgH) repertoire, we analyzed tet-LIN28B mice on the B1-8hi BCR transgenic background, which expresses a pre-rearranged, non-self-reactive IgH chain that is specific to the hapten 4-hydroxy-3-nitrophenylacetyl (NP) when paired with the Igλ light chain(*53*). Strikingly, the emergence of both overall Igλ⁺ and NP-specific immB cells was significantly reduced by LIN28B on the B1-8hi background, to levels comparable to those observed in neonatal mice (Figure 4A-B). This result suggests that LIN28B suppresses secondary rearrangements independently of IgH repertoire composition. In addition, LIN28B still resulted in reduced surface CD25 levels and a mild decrease in the representation of the small pre-B cell subset in the context of a fixed IgH chain (Figure 4C, Supplementary Figure 4B). These findings cannot be explained by differential usage of the transgenic B1-8hi IgH(*54*) (Supplementary Figure 4C) and thus uncouple LIN28B-driven phenotypes from alterations in BCR self-reactivity. Moreover, the LIN28B dependent suppression of CD25 surface expression was retained on the SLC–/– background (Figure 4D), demonstrating that LIN28B interferes with the CD25⁺ pre-B_REC_ state independently of pre-BCR or mature BCR signaling.

**Figure 4.**
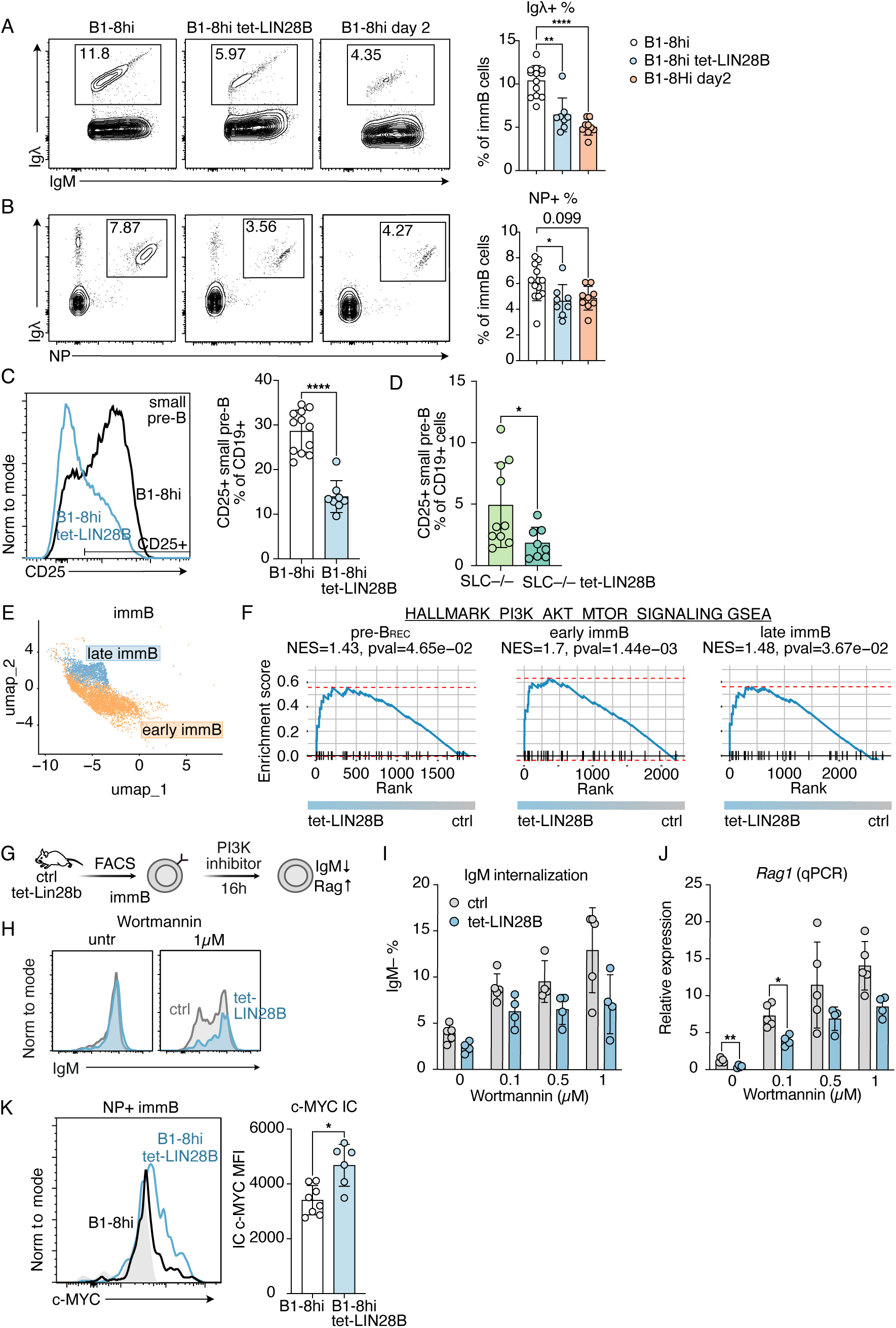
LIN28B suppresses secondary rearrangements in immB cells independently of IgH specificity. A. Representative FACS plot and percentages of Igλ⁺ cells out of immB cells in adult B1-8hi (n=13), B1-8hi tet-LIN28B (n=8), and neonate B1-8hi (postnatal day 2, n=9). Mean ±SD. Kruskal-Wallis test, with Dunn’s correction. B. Representative FACS plot and percentages of 4-hydroxy-3-nitrophenylacetyl (NP)-reactive (NP⁺Igλ⁺) cells out of immB cells in adult B1-8hi (n=13), B1-8hi tet-LIN28B (n=8), and neonate B1-8hi (postnatal day 2, n=9). Mean ±SD. Kruskal-Wallis test, with Dunn’s correction. C. Representative FACS histogram of CD25 surface expression on small pre-B cells and percentages of CD25⁺ small pre-B cells out of CD19⁺ cells in B1-8hi (n=12) and B1-8hi tet-LIN28B (n=8) mice. Mean ±SD. Mann-Whitney test. D. Percentages of CD25⁺ small pre-B cells out of CD19⁺ cells in Surrogate Light Chain-deficient (SLC–/–, n=10) and SLC–/– tet-LIN28B (n=8) mice. Mean ±SD. Mann-Whitney test. E. Re-clustering of immB cells from scRNAseq data in Figure 1B. F. Gene Set Enrichment Analysis (GSEA) in tet-LIN28B vs ctrl pre-B_REC_, early immB and late immB cells for the PI3K/AKT/mTOR signaling pathway Hallmark Geneset from the Mouse Molecular Signatures Database (MSigDB)(*69*). G. Experimental layout of *in vitro* PI3K inhibition: immB cells were FACS sorted and cultured *in vitro* in presence of increasing concentrations of Wortmannin, and evidence of back-differentiation was measured by IgM downregulation by flow cytometry and *Rag1* induction by RT-qPCR. H. Representative FACS histogram of surface IgM expression of immB cells sorted from ctrl or tet-LIN28B mice after 16 hours with or without (untreated, untr) Wortmannin (1 µM). I. Percentage of IgM^−^ cells in overnight culture of immB cells from ctrl (n=5) or tet-LIN28B (n=4) mice treated with the indicated concentrations of Wortmannin. Mean ±SD. Two-way ANOVA test, with Šídák correction. J. *Rag1* expression assessed by qRT-PCR in cells in I. Expression is normalized to *β-Actin* and presented as relative to the average of the untreated ctrl samples from the same experiment. Mean ±SD. Two-way ANOVA test, with Šídák correction. K. Representative FACS histogram and median fluorescence intensity (MFI) of c-MYC intracellular (IC) expression on immB cells in B1-8hi (n=8) and B1-8hi tet-LIN28B (n=6) mice. Mean ±SD. Mann-Whitney test. Only significant comparisons shown, p/adj.p: * < 0.05, ** < 0.01, *** < 0.001, **** < 0.0001.

We next investigated the BCR specificity independent mechanism underlying the effects of LIN28B on late pre-B_REC_ cells. Signaling through the Phosphoinositide 3-kinase (PI3K) pathway is a powerful regulator of the balance between receptor editing and positive selection. Its interruption triggers back-differentiation of immB cells to a pre-B_REC_-like stage to undergo secondary rearrangements while its positive regulation can promote developmental progression even in the face of self-reactive BCRs(*29, 31, 51, 55*). We have previously demonstrated that LIN28B augments PI3K/AKT/mTOR signaling in immB cells (*37*) and this was supported by our scRNAseq data (Supplementary Figure 4D, Supplementary Table 1). To trace when this effect emerges, we re-clustered immB cells into early and late subsets (Figure 4E, Supplementary Figure 4E). Gene set enrichment analysis showed that the PI3K/AKT/mTOR signature became significantly enriched as LIN28B-expressing cells transitioned from pre-B_REC_ into early immB cells, the juncture at which cells actively choose between the forward and backward differentiation trajectories (Figure 4F, Supplementary Table 1). To test whether the LIN28B-driven reduction in the secondary rearrangement is dependent on PI3K, we FACS sorted tet-LIN28B and ctrl immB cells and cultured them in the presence or absence of the PI3K inhibitor Wortmannin to mimic attenuated tonic signaling as previously described (*51*). At 16 hours post treatment, Wortmannin triggered *Rag1* reactivation and surface IgM downregulation in both genotypes in a dose dependent manner, confirming that the secondary rearrangement machinery remains PI3K sensitive in tet-LIN28B cells (Figure 4G-J, Supplementary Figure 4F). Interestingly, LIN28B-expressing cells never reached control levels in either readout, consistent with an elevated basal PI3K/AKT/mTOR activity that favors developmental progression over secondary rearrangements.

MYC operates in a positive feedback loop with PI3K in B cells and is a known driver of positive selection (*56*). Extending our analysis to MYC, we found that NP-restricted B1-8hi immB cells showed a modest elevation of MYC protein levels upon LIN28B expression (Figure 4K). These data critically extend previous findings showing that LIN28B amplifies the PI3K/MYC feedback loop and promotes positive selection (*37*) by uncoupling these effects from BCR repertoire composition. Taken together, LIN28B acts through a BCR specificity-agnostic mechanism to bias developing B cells towards selection over editing, providing a potential explanation for the retention of self-reactive clones early in life.

### CD25 marks a pre-B_REC_ cell state that is metabolically quiescent and rearrangement-permissive

To investigate the transcriptional state associated with pre-B_REC_ CD25 expression, we re-analyzed a published BCP scRNAseq dataset in which pre-B cells were sorted based on CD25 status (*45*). Although individual transcript differences were modest (Supplementary Table 3), CD25^−^ pre-B_REC_ cells were significantly enriched for E2F and MYC targets as well as oxidative phosphorylation gene signatures, linking CD25 expression to a proliferatively and metabolically more quiescent state (Figure 5A-C, Supplementary Figure 4G). This finding was corroborated by our own whole transcriptome analysis (WTA) scRNAseq data from control mice, which showed that transcripts negatively correlated with *Il2ra* were most significantly enriched for MYC and E2F targets (Figure 5D, Supplementary Table 4). In line with this, we found a mild but consistent decrease in MYC protein levels in CD25⁺ compared to CD25^−^ pre-B cells as measured by intracellular staining both on IgH polyclonal and B1-8hi transgenic backgrounds (Figure 5E-F). Together, these findings identify CD25 as a marker of the quiescent state that characterizes adult pre-B_REC_ cells, consistent with LIN28B suppressing CD25 expression early in life while sustaining an elevated MYC-driven anabolic program (*37*).

**Figure 5.**
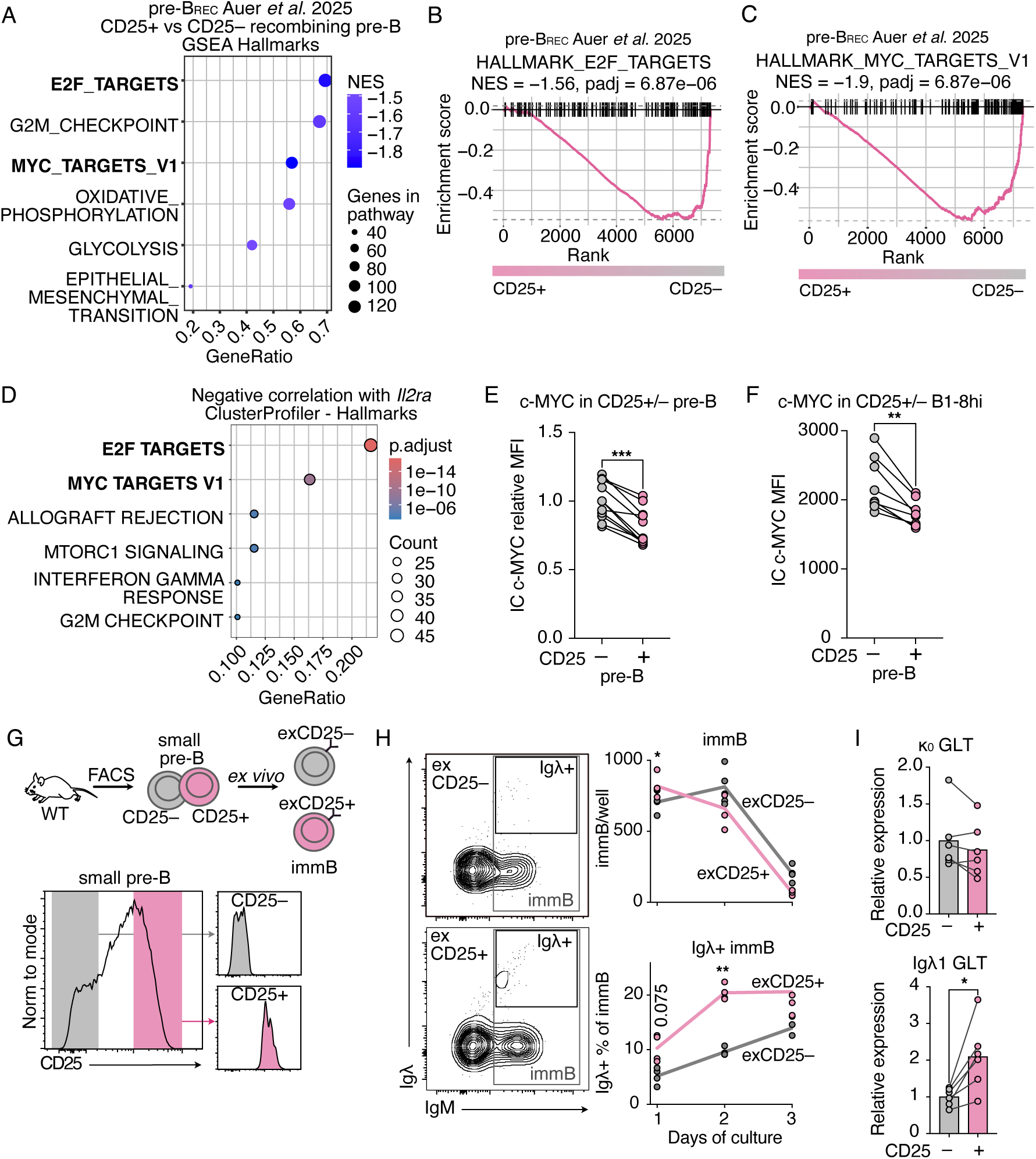
CD25 marks a metabolically restricted pre-B cell state biased for receptor editing. A. GSEA in CD25⁺ vs CD25^−^ pre-B_REC_ identified by reanalyzing a published dataset (*45*) for Hallmark Genesets of the MSigDB(*69*). The dot plot shows significant genesets negatively associated with CD25⁺ pre-B_REC_. NES=Normalized Enrichment Score. B. Enrichment plot for the E2F target geneset in A. C. Enrichment plot for the MYC target geneset in A. D. Enrichment for Hallmark Genesets of the MSigDB(*69*) assessed using ClusterProfiler in genes negatively correlated with *Il2ra* expression in the scRNAseq data from ctrl mice (Spearman correlation r < -0.1). E. MFI of c-MYC IC staining in CD25⁺ and CD25^−^ pre-B cells from WT mice, presented as relative to the mean of CD25^−^ cells from the same experiment. Connecting lines link CD25⁺ and CD25^−^ cells from the same mouse. Two-tailed Wilcoxon matched-pairs signed rank test. F. MFI of c-MYC IC staining in CD25⁺ and CD25^−^ pre-B cells in B1-8hi mice. Connecting lines link CD25⁺ and CD25^−^ cells from the same mouse. Wilcoxon matched-pairs signed rank test. G. Experimental layout: small pre-B cells were FACS sorted according to CD25 status and cultured with 0.05 ng/ml murine IL-7 (mIL-7) to induce differentiation into immB cells, detected by flow cytometry. H. Representative FACS plot at day 2 post culture, immB cell counts, and Igλ⁺ percentage of immB cells differentiated from sorted CD25⁺ (exCD25⁺, n=4) and CD25^−^ (exCD25^−^, n=4) small pre-B cells cultured as described in G. Two-way ANOVA test, with Šídák correction. I. *Igκ_0_* and *Igλ1* GLT expression in sorted CD25⁺ (n=6) and CD25^−^ (n=6) small pre-B cells assessed by RT-qPCR. Two-tailed Wilcoxon matched-pairs signed rank test. Only significant comparisons shown, p/adj.p: * < 0.05, ** < 0.01, *** < 0.001, **** < 0.0001.

Since RAG-mediated recombination is associated with a quiescent metabolic state (*24*), we hypothesized that CD25 expression levels might reflect a broader cellular state linked to the capacity to undergo secondary rearrangements. To test this, we sorted CD25⁺ and CD25^−^ small pre-B cells from wildtype (WT) mice and allowed their spontaneous differentiation into immB cells in culture (Figure 5G). CD25⁺ sorted small pre-B cells generated a higher proportion of Igλ⁺ immB compared to their CD25^−^ sorted counterparts (Figure 5H) and displayed higher expression of *Igλ* but not *Igκ* germline transcripts (Figure 5I). We conclude that CD25 marks a pre-B cell state more permissive to sustaining extended secondary IgL rearrangements.

### LIN28B expression in early life accelerates B cell maturation and bone marrow output

To determine whether LIN28B-driven positive selection translates into accelerated BM B cell output, we first assessed the kinetics of pro-B to immB cell maturation *in* vitro. Pro-B cells were FACS sorted and allowed to differentiate in a stroma-free culture under low IL-7 conditions (Figure 6A). Accelerated immB differentiation was observed for tet-LIN28B pro-B cells (Figure 6B) and was phenocopied by neonatal pro-B cells in an endogenous LIN28B-dose-dependent fashion (Figure 6C). Adding to this, scRNAseq data showed that *Cxcr4* was downregulated in both LIN28B-expressing adult and neonatal pre-B_REC_ and immB cells. (Figure 6D-F). CXCR4 is a defining chemokine receptor for pre-B cells, retaining them in the bone marrow niche to complete IgL recombination. Its expression is suppressed by positive selection or under conditions of elevated PI3K signaling (*29, 30*) and upregulated by self-reactive BCR engagement(*30, 52, 57*). Surface CXCR4 expression was mildly decreased, though not significantly, in freshly isolated tet-LIN28B immB cells, likely reflecting rapid BM egress of low expressors (Figure 6G). To circumvent this capture issue, we assessed CXCR4 surface expression following *in vitro* differentiation of sorted pro-B cells into immB cells. Indeed, this approach revealed significantly decreased CXCR4 surface expression (Figure 6H-I). Together, these data show that LIN28B not only accelerates B cell maturation but also promotes BCP bone marrow egress.

**Figure 6.**
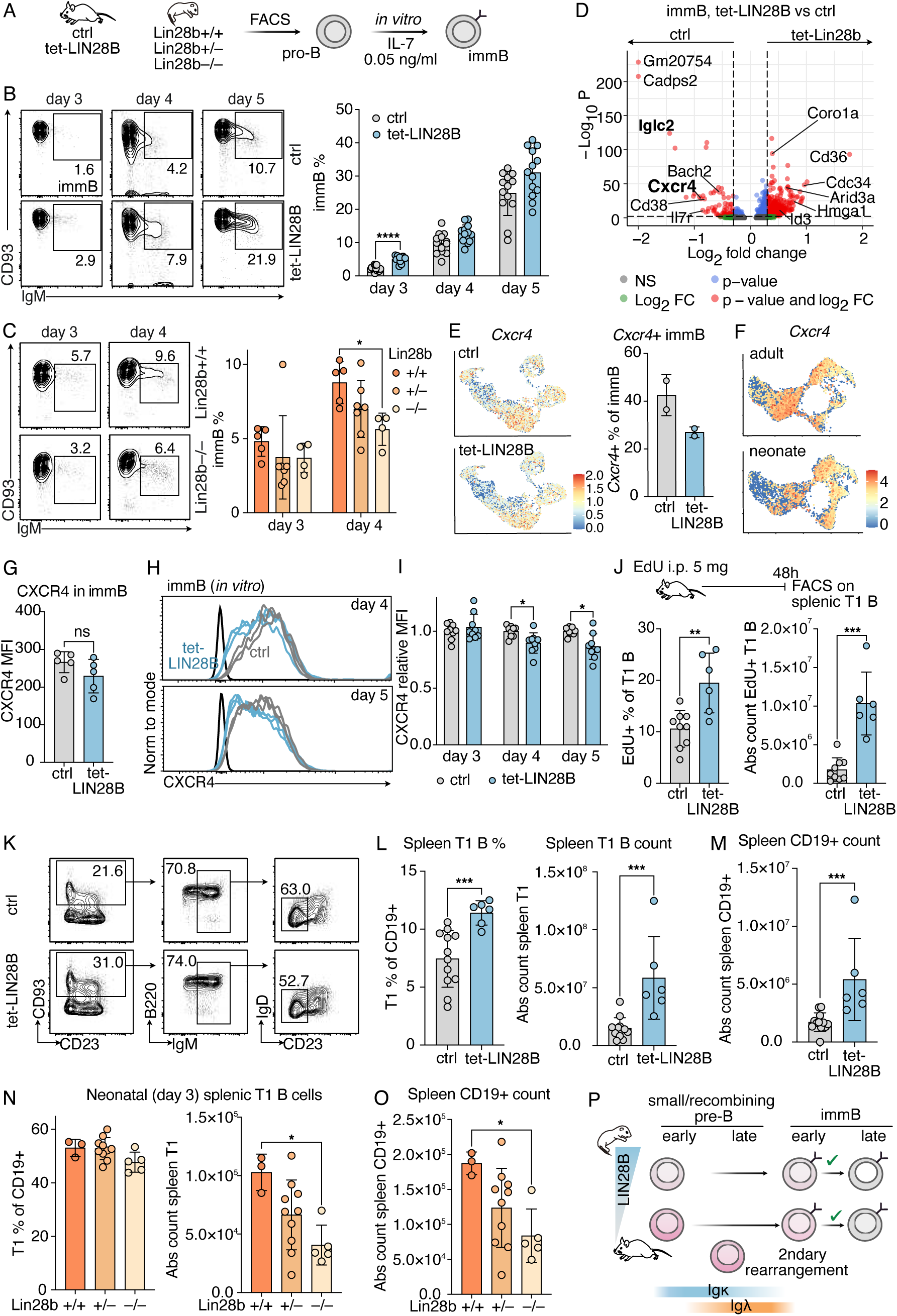
LIN28B accelerates bone marrow B cell differentiation and output in early life. A. Experimental layout: pro-B cells from ctrl and tet-LIN28B adult mice, or from Lin28b+/+, +/–, and –/– neonate mice were FACS sorted and cultured *in vitro* with 0.05 ng/ml mIL-7 to induce differentiation into immB cells, detected by flow cytometry. B. Representative FACS plots and percentages of immB cells differentiated from sorted pro-B cells from adult ctrl (n=13) and tet-LIN28B (n=13) mice. Mean ±SD. Two-way ANOVA test, with Šídák correction. C. Representative FACS plots and percentages of immB cells differentiated from sorted pro-B cells from postnatal day 2-3 neonates *Lin28b*+/+ (n=5), +/– (n=7) and –/– (n=4) mice. Mean ±SD. Two-way ANOVA test, with Tukey correction for multiple comparisons. D. Volcano plot of differentially expressed genes in tet-LIN28B vs ctrl immB cells from the scRNAseq dataset. E. *Cxcr4* mRNA expression ctrl and tet-LIN28B cells from the WTA scRNAseq experiment and percentages of ctrl and tet-LIN28B immB cells where *Cxcr4* expression was detected. Mean ±SD. F. *Cxcr4* mRNA expression ctrl adult and neonate cells from the targeted scRNAseq experiment. G. CXCR4 MFI in freshly isolated ctrl and tet-LIN28B immB cells measured by flow cytometry. Mean ±SD. Mann-Whitney test. H. Representative FACS histogram overlays of CXCR4 surface expression in immB cells from ctrl and tet-LIN28B mice differentiated *in vitro* for 4-5 days from sorted pro-B as described in A. Black line represents FMO. I. CXCR4 MFI in immB cells from ctrl (n=9) and tet-LIN28B (n=9) mice differentiated in culture for 3 to 5 days from sorted pro-B as described in A. Values presented as relative to the mean of ctrl samples from the same experiment. Two-way ANOVA test, with Šídák correction. J. EdU pulse chase experiment: mice were injected intraperitoneally (i.p.) with a single dose of 5 mg of EdU and labelling was assessed in spleen transitional 1 (T1) B cells after 48h. The percentages and absolute counts of EdU⁺ cells in splenic T1 B cells of ctrl (n=9) and tet-LIN28B (n=6) mice are shown in bar graphs. Mean ±SD. Mann-Whitney test. K. Gating strategy from CD19⁺ cells and representative FACS plot of T1 B cells in the spleen of ctrl and tet-LIN28B adult mice. L. Percentages and absolute counts of T1 B cells in the spleen of ctrl (n=12) and tet-LIN28B (n=6) adult mice. Mean ±SD. Mann-Whitney test. M. Absolute numbers of overall CD19⁺ B cells in the spleen of ctrl (n=12) and tet-LIN28B (n=6) adult mice. Mean ±SD. Mann-Whitney test. N. Percentages and absolute counts of T1 B cells in the spleen of neonate Lin28b+/+ (n=3), +/– (n=10) and –/– (n=5) mice. Mice of different genotypes are littermates from three litters. Mean ±SD. Kruskal-Wallis test, with Dunn’s correction. O. Absolute counts of overall CD19⁺ B cells in the spleen of neonate Lin28b+/+ (n=3), +/– (n=10) and –/– (n=5) mice. Mice of different genotypes are littermates from three litters. Mean ±SD. Kruskal-Wallis test, with Dunn’s correction. Only significant comparisons shown, p/adj.p: * < 0.05, ** < 0.01, *** < 0.001, **** < 0.0001. P. Working model.

To resolve whether this led to accelerated BM B cell output, we administered intraperitoneally a single pulse of 5-Ethynyl-2′-deoxyuridine (EdU, 5 mg per mouse) and tracked labeling in splenic transitional 1 (T1) B cells 48 hours later. Tet-LIN28B mice showed a significant increase in EdU⁺ T1 B cells compared to ctrl indicating an accelerated BM to spleen transit (Figure 6J), in agreement with our previous observations (*37*). Consistent with LIN28B enhancing B cell output, we observed an increase in splenic transitional 1 (T1) B cells (Figure 6K-L) and an overall increase in splenic B cell numbers (Figure 6M) upon LIN28B induction. Similarly, T1 B cell numbers were higher in WT neonates compared to their Lin28b–/– counterparts (Figure 6N), resulting in higher overall CD19⁺ B cell counts in neonates at 3 days of age (Figure 6O). Taken together, these findings establish that LIN28B accelerates B cell maturation and bone marrow output at the expense of stringent receptor editing, promoting an early-life program that prioritizes the rapid seeding of B cells and permissive to retaining of a wider range of self-reactive BCR specificities (Figure 6P).

## DISCUSSION

B cells developing during fetal and neonatal life exhibit enhanced self-reactivity in mouse and human compared to their adult counterparts (*4, 6, 7*), indicating altered regulation of repertoire establishment. To dissect changes in tolerance mechanisms during ontogeny, we took advantage of tet-LIN28B inducible mice, which recapitulate key features of early-life B lymphopoiesis(*35, 37*). Using scRNAseq of BM B cell progenitors, we uncovered a decreased abundance of pre-B_REC_ cells, undergoing IgL recombination, in LIN28B expressing and neonatal cells. This decrease reflects a reduced window for secondary rearrangements. While previous work established that LIN28B augments positive selection of self-reactive CD5⁺, Nur77⁺ immB cells in early life (*37, 38*), the present study uncovers regulation at the level of IgL secondary rearrangements and, importantly, in a manner that is uncoupled from strong self-antigen engagement. Furthermore, we identified an adult enriched pre-B cell state that is marked by CD25 expression and a transcriptomic signature of metabolic quiescence. Despite being biased towards receptor editing, this state can be dissociated from both pre-BCR signaling strength and IgH specificity and is effectively blunted by LIN28B. These findings reveal that an early-life developmental program can modulate the BCR-instructed checkpoint of central B cell tolerance, increasing its permissiveness in a specificity-agnostic manner. Taken together, these insights highlight central tolerance stringency as a developmentally controlled threshold and provide an experimental platform for its deliberate manipulation.

The best characterized function of LIN28B, conserved from *C. elegans* to human, is the post-transcriptional suppression of let-7 microRNA biogenesis (*41*), which in turn relieves the repression of let-7 target genes regulating cell expansion and metabolism. Among these, let-7 directly represses multiple components of the PI3K/AKT/mTOR pathway as well as *Myc* itself(*58, 59*), providing a likely basis for the elevated PI3K and MYC activity signatures observed in LIN28B-expressing pre-B_REC_ cells. Consistent with the broader mode of microRNA-mediated gene regulation, the observed effects of tet-LIN28B were relatively mild. However, the acceleration and qualitative shift in B cell output suggest that such individual fine-tuning events are consequential once combined. Thus, given that elevated PI3K activity, driven in part by MYC(*56*), is known to suppress RAG1/2-mediated recombination as well as CXCR4 expression, the let-7 dependent mechanism is sufficient to explain the reduction of receptor editing and premature dissolution of pre-B_REC_ identity (*30, 31*). Beyond let-7 suppression, LIN28B may additionally reinforce this anabolic state through directly binding ribosomal protein transcripts to promote ribosome biogenesis, further amplifying the biosynthetic capacity of fetal-like pre-B cells(*35*). The precise contribution of each of these mechanisms, and how they are coordinated to enforce the receptor editing checkpoint, remains to be defined.

While CD25 has long been used as a marker for pre-B cell staging, its expression is not ubiquitous among pre-B cells, and its functional significance has remained elusive. A recent study highlighted how CD25 is expressed predominantly in recombining pre-B cells as opposed to cycling pre-B cells(*45*). Our results add another layer of resolution, showing that CD25 expression level is associated with the propensity of pre-B_REC_ to engage in secondary rearrangements, with CD25^high^cells more likely to emerge as Igλ⁺ immB compared to CD25^low^ cells. Importantly, this stratification is linked to differences in MYC activity and metabolic signatures and operates independently of BCR specificity. We therefore propose that CD25 expression at the pre-B_REC_ stage serves as a functional marker for metabolically restricted cells primed to engage in receptor editing. Whether CD25 is itself functionally involved in regulating editing propensity remains an important open question, as a recent genetic study failed to find functional consequences of CD25 deficiency during B cell development(*60*). Our findings shed new light on the functional significance of CD25 during B cell development, establishing it as a broadly applicable parameter for the study of receptor editing propensity. At the same time, caution should be exercised when using CD25 as a pre-B cell marker in early life, where LIN28B-mediated suppression of the CD25^high^ state confounds its interpretation.

The extent to which LIN28B affects clonal deletion or peripheral tolerance was not addressed in this study and remains an open question. Previous work showed that receptor editing response is already active in early life when LIN28B is endogenously expressed (*7, 33*), suggesting that the most self-reactive specificities are likely still purged, leaving only moderately self-reactive BCRs to progress into the periphery. Nevertheless, previous observations by Cornall and colleagues demonstrate that LIN28B can restore positive selection of self-reactive B cells in a BCR transgenic model (*38*), though whether this reflects effects on receptor editing, clonal deletion, or peripheral survival competition was not resolved in that study. Work modeling constitutive PI3K activation found that although central tolerance was breached, self-reactive B cells still failed to secrete autoantibodies *in vivo*(*29*), suggesting that late peripheral checkpoints may further offset the risk incurred by a more relaxed central tolerance threshold.

When considering the evolutionary value of a relaxed early-life tolerance threshold, it is important to note that while stringent receptor editing purges potentially harmful self-reactivity, it also risks depleting beneficial BCR clonotypes with reactivity to shared conserved host structures and foreign antigens. Such useful self-reactivity has been described as “autoreactive by design” as they derive from a largely germline encoded repertoire and include innate-like B-1 cells directed against phospholipid moieties involved in tissue homeostasis and early defense(*8*). This trade-off between the potential dangers of self-reactivity and benefits of broad protection may be particularly consequential during the neonatal period, when the immune system must rapidly establish a functional repertoire in the absence of immunological memory. Notably, this is a developmental window in which the inherently restricted diversity of the early-life repertoire (*61, 62*) may itself limit the potential for pathogenic self-reactivity, making relaxed central tolerance more defensible. From this perspective, the action of LIN28B may represent a selected feature rather than a flaw of early-life B cell development, intended to preserve certain useful self-reactive specificities. Analogous observations have been made in the T cell lineage, where a similar bias toward innate-like effector fates and accelerated maturation has been described during early ontogeny(*63, 64*). Interestingly, both B-1 cells and regulatory T cells of early-life origin share immunoregulatory effector functions and self-reactive features. Together, these parallels suggest a shared adaptive paradigm of the fetal-neonatal immune system, optimized for rapid colonization of peripheral niches and the incorporation of unique functionalities into the long-lived lymphocyte pool to promote tissue homeostasis.

## MATERIALS AND METHODS

### Study design

The aim of the study was to investigate differences between early life and adult B cell differentiation at steady state, and to determine the contribution of LIN28B to neonatal-specific features. An unbiased approach informed subsequent hypothesis and interpretation. B cell differentiation was compared between wildtype neonates (2-3 days after birth) and adults (8-22 weeks of age). The role of LIN28B was investigated using two complementary models: adult mice with doxycycline-induced ectopic LIN28B expression and Lin28b–/– neonates. Littermates were used as controls, and investigators were not blinded to the genotypes.

B cell progenitors were studied by scRNAseq, flow cytometry, (qRT)-PCR and *ex vivo* culture. For scRNAseq, two samples per group were included, with each neonatal sample consisting of a pool of 8-9 pups from the same litter. All other experiments included at least three biological replicates from at least two independent experiments and litters. Outliers were retained unless there was evidence of a technical error. Statistical tests used nonparametric test and p-values were corrected for multiple testing when more than two groups were compared.

### Mice

B6.Cg-*Col1a1*^tm2^ *^(tetO-LIN28B)Gqda^*/J mice (Jax stock #023911) were obtained from the laboratory of Dr. George Daley(*58*) (Harvard Medical School). B6-Cg-*Gt(ROSA)26Sor^tm1(rtTA*M2)Jae^*/J mice (Jax stock #006965) were from The Jackson Laboratory(*65*). These strains were intercrossed to obtain trans-heterozygous tet-LIN28B mice. LIN28B expression was induced by feeding doxycycline-containing chow (200 mg/kg, SSniff, #A112D70203) for at least 10 days. Doxycycline-treated littermate harboring the *R26*^m2rtTA^ allele were used as controls. *Lin28b–/–* (B6.Cg-*Lin28b^tm1.1Gqda^/J* Jax #023917) mice were from The Jackson Laboratory(*66*). SLC–/– mice (*40*) were obtained from the laboratory of Lill Mårtensson and crossed with tet-LIN28B mice. B1-8hi mice (Jax stock #007594) were obtained from the laboratory of Taras Kreslavskiy and crossed with tet-LIN28B mice. Mice used in B1-8hi experiments were heterozygous for the B1-8hi and CD45.1/2 alleles. For experiments involving SLC–/– or B1-8hi mice, doxycycline-treated littermates heterozygous for either the *R26*^m2rtTA^ or the *Col1a1*^tm2^ *^(tetO-LIN28B)Gqda^* allele served as controls. All adult mice were between 8 and 22 weeks old. Neonate mice were 2-3 days old. All animal procedures were performed, and mice were housed, in accordance with ethical permits approved by the Swedish Board of Agriculture.

### Flow cytometry analysis and sorting

Bone marrow (BM) cells were extracted by crushing bones with mortar and pestle as previously described(*67*). Adult BM was subjected to red blood cell lysis using Ammonium-Chloride-Potassium (ACK) Lysing Buffer (Gibco, #A1049201). For B cell progenitors (BCP) isolation, lineage-positive cells (Ter119^+^Gr1^+^CD3^+^) were depleted by magnetic-activated cell sorting (MACS; Miltenyi Biotec) according to the manufacturer’s instructions. All steps were performed with HBSS (Gibco, #14175129) supplemented with 0.5% BSA (Sigma-Aldrich, #A7030) and 2 mM EDTA (Invitrogen, #15575020). Cells were stained with antibodies at a density of 1-5 × 10^6^ cells/100 μL volume for 30 min in the dark at 4°C. Viability was assessed using 7-Amino-Actinomycin D (7-AAD, Sigma-Aldrich, #SML1633). Flow cytometry experiments were performed at the Lund Stem Cell Center FACS Core Facility (Lund University) on BD FACS Symphony S6, BD FACS Aria III, BD Fortessa, BD Fortessa X20 and BD Symphony A1 instruments. Populations were sorted using a 70-or 85-µm nozzle, a “0.32.0” precision mask, and a maximum event rate of 5,000 events/s. Populations were defined as follows:

- pro-B: Ter119^−^CD3^−^Gr1^−^CD19⁺B220⁺CD93⁺IgM^−^CD117⁺

- pre-B: Ter119^−^CD3^−^Gr1^−^CD19⁺B220⁺CD93⁺IgM^−^CD117^−^CD25⁺/^−^FSC-A^high/low^

- immB: Ter119^−^CD3^−^Gr1^−^CD19⁺B220⁺CD93⁺IgM⁺

For intracellular staining, cells were stained for surface markers and then fixed and permeabilized using the Foxp3/Transcription Factor Fixation/Permeabilization kit (eBioscience, #00-5521-00) before staining with intracellular markers. For c-MYC detection, cells were stained with a primary antibody, blocked with goat serum (Abcam, #ab7481), and subsequently stained with a secondary F(ab’)₂ goat anti-rabbit IgG (H+L) antibody. Flow cytometry data were analyzed using FlowJo v10.10.1.

### Single cell RNA sequencing WTA: library preparation and sequencing

BM BCPs (Ter119^−^CD3^−^Gr1^−^CD19⁺B220⁺CD93⁺) and recirculating B cells (Ter119^−^CD3^−^Gr1^−^ CD19⁺B220⁺CD93^−^) were FACS sorted and pooled at a 90:10 ratio. Cells were labeled using the BD Mouse immune Single-cell Multiplexing kit (BD, #633793), and cell number and viability were measured by Trypan blue staining on TC20 Automated Cell Counter (BioRad). A total of 60 000 cells were loaded onto a BD Rhapsody Cartridge (BD, #666262) using the BD Rhapsody Express Instrument. Single cell bead capture, cell lysis and barcoding and cDNA synthesis were performed according to the manufacturer’s instructions. scRNAseq cDNA libraries were purified with AMPure XP Reagent (Beckman coulter, #A63880), quantified with a Qubit Fluorometer using dsDNA HS Assay Kit (Fisher Scientific, #Q32851), and quality controlled on the Agilent 2100 Bioanalyzer using the High Sensitivity DNA Kit (Agilent, #5067-4626). Final libraries were sequenced on Illumina NovaSeq 6000 sequencing system using the Nova seq 6000 kit S2 v1.5 100 cycles. (Illumina, #20028318) at the Center for Translational Genomics (CTG), Lund University.

### Single cell RNA sequencing WTA: data preprocessing and analysis

FASTQ files were processed with the Seven Bridges cloud-based platform using the BD Rhapsody Sequence Analysis Pipeline, to generate a count matrix. Downstream analyses were performed in Jupyter notebooks using Seurat v5.1.0 in R v4.4.0 (R Core Team; www.R-project.org/).

Undetermined and multiplet data were excluded. Cells passing the following quality-control criteria were retained: 1200 < nFeature_RNA < 6000, nCount_RNA < 25000, and 5–13% mitochondrial transcripts. Data were normalized and the highly variable features were identified after excluding BCR genes, as previously described (*68*), and mitochondrial transcripts. Cell-cycle scores were calculated using “CellCycleScoring” based on Seurat-provided G2/M-and S-phase gene sets converted to their mouse orthologs, followed by regression using “SCTransform”. BCR and mitochondrial genes were subsequently excluded from SCT variable features. Clustering was performed according to the Seurat workflow (https://satijalab.org/seurat/articles/pbmc3k_tutorial.html) using 20 dimensions in “FindNeighbors”, followed by “FindClusters” at a resolution of 0.2. Differentially expressed genes (DEGs) between control and tet-LIN28B cells were determined using Seurat’s “FindMarkers” function. Pseudotime analysis was performed using Monocle3 v1.3.7. The pre-B^REC^ and immB clusters were separately re-clustered using FindClusters at resolutions of 0.2 and 0.1, respectively. Volcano plots of DEGs were generated using EnhancedVolcano v1.28.2 in R v4.5.3. Let-7 target enrichment was calculated using fgsea v1.36.2 and the “LET_7A_5P_LET_7C_5P_LET_7E_5P_MIR_98_5P” geneset from the Mouse Molecular Signatures Database (MSigDB)(*69*) v26.1.0 package in R v4.5.3. Hallmark term enrichment was performed using the “enricher” function in ClusterProfiler v4.18 on DEGs with adjusted p-value < 0.05 and log2FC > 0 and genes negatively correlated with *Il2ra* expression in control cells (Spearman correlation < –0.1).

### Targeted single cell RNA sequencing: library preparation and sequencing

BM BCPs (Ter119^−^CD3^−^Gr1^−^CD19⁺B220⁺CD93⁺) were labeled using the BD Mouse immune Single-cell Multiplexing kit (BD, #633793) according to manufacturer instructions, FACS sorted and pooled. The pooled cells were stained with the following BD oligonucleotide-conjugated antibodies: Ms CD19 Oligo AMM2007 1D3 (#940111), Ms IgM Oligo AMM2031 II/41 (#940135), Ms CD117 (c-Kit) Oligo AM (#940127), Ms CD25 Oligo AMM2164 3C7 25Tst (#940356), Ms CD5 Oligo AMM2043 53-7 (#940147). Cell number and viability were measured by Calcein AM (Thermo Fisher Scientific, #C1430) and DRAQ7™ (BD Biosciences, #564904) staining using a BD Rhapsody Scanner. A total of 50 000 cells were loaded onto a BD Rhapsody Cartridge (BD, #633733) using the BD Rhapsody Express Instrument. Single cell bead capture, cell lysis and barcoding, and cDNA synthesis were performed according to manufacturer’s instructions. For targeted amplification, the BD Rhapsody Immune Response Panel Mm (BD, #633753) was supplemented with custom primers to detect a total of 463 transcripts (*47*) (Supplementary Table 2). scRNAseq cDNA libraries were purified with AMPure XP Reagent (Beckman coulter, #A63880), quantified with a Qubit Fluorometer using dsDNA HS Assay Kit (Fisher Scientific, #Q32851), and quality controlled on an Agilent 2100 Bioanalyzer using the High Sensitivity DNA Kit (Agilent, #5067-4626). Final libraries were sequenced on an Illumina NextSeq using the 500/550 High output kit v2.5 (150 cycles).

### Targeted single cell RNA sequencing: data preprocessing and analysis

FASTQ files were processed on the Seven Bridges cloud-based platform using the BD Rhapsody Sequence Analysis Pipeline, to generate count matrices. Downstream analyses were performed with Seurat v5.5.0 in R v4.5.3. Undetermined cells and multiplets were excluded, and cells passing the following quality-control criteria were retained: nFeature_RNA > 40, nFeature_RNA < 150, nCount_RNA < 1900. Clustering was performed according to the Seurat workflow using 20 dimensions in “FindNeighbors” function, followed by “FindClusters” at a resolution of 0.5.

### Analysis of published scRNAseq data

Published scRNAseq data from wildtype cells in (*45*) were reanalyzed using Seurat package v5.5.0 in R v4.5.3. Cells were retained based on the following quality-control criteria: nFeature_RNA > 200, nFeature_RNA < 5000, nCount_RNA > 1500, and mitochondrial transcripts < 5%. Clustering was performed using “FindNeighbors” with 30 dimensions, followed by “FindClusters” at a resolution of 0.3. Within the *Rag1*-expressing pre-B cluster (here termed pre-B^REC^), DEGs between CD25⁺ and CD25⁻ pre-B cells, corresponding to PreBII and PreBI cells, respectively, in the original publication, were identified using Seurat’s “FindMarkers” function. Gene set enrichment analysis was performed on DEGs using “fgsea” and Hallmark gene sets from the MSigDB (*69*).

### RT-qPCR

RNA was extracted from sorted cells stored in DNA/RNA shield (Zymo Research, #R1100) or RNAzol RT (Sigma-Aldrich, #R4533) using the Quick-DNA/RNA Microprep Plus Kit (Zymo Research, #D7005) or Directzol Micro RNA prep (Zymo Research, #R2060), respectively. RNA was reverse-transcribed using TaqMan® Reverse Transcription Reagents (Fisher scientific, #N8080234) with random primers.

IgL transcripts were measured using the QuantiTect SYBR Green PCR Kit (Qiagen, #204143) with an annealing temperature of 60°C. Rearranged *Igκ* transcripts were amplified using a degenerate *Vκ* primer and a *Jκ1* primer were used. *B2m* was used as reference gene. Primer sequences are listed in Supplementary Table 5.

*Rag1* was measured using KAPA Probe Fast qPCR Master Mix (Sigma-Aldrich, #KK4706) with an annealing temperature of 60°C, and *Actb* as reference gene. The following probes from IDT were used: *Rag1* Mm.PT.58.6440291, *ActB* Mm.PT.39a.22214843.g.

### *Igκ* rearrangement analysis

DNA was extracted from sorted cells stored in DNA/RNA shield (Zymo Research, #R1100) using the Quick-DNA/RNA Microprep Plus Kit (Zymo Research, #D7005). *VκJκ* joints were amplified for 32 cycles using Platinum SuperFi II DNA Polymerase (ThermoFisher, #12361010), a degenerate *Vκ* primer, and a *Jκ5* primer, as previously described (*32*) (Supplementary Table 5). *Actb* was amplified as a loading control. PCR products were separated by electrophoresis on 1.5% agarose gels and imaged using a ChemiDoc (Bio-Rad). Band intensities were quantified using ImageJ.

### *Ex vivo* culture assays

Small pre-B (CD25⁺ or CD25^−^) or pro-B cells were FACS sorted and cultured in B cell medium consisting of RPMI 1640 (Gibco, #11875093), 10% Fetal Bovine Serum (Gibco, #26400044), 2% HEPES (ThermoFisher Scientific, #15630080), 1% MEM non-essential amino acids (ThermoFisher Scientific, #11140050), 1% sodium pyruvate (ThermoFisher Scientific, #11360070), 1% P/S (HyClone, #SV30010), 0.2% β-mercaptoethanol (ThermoFisher Scientific, #31350010), supplemented with 0.05 ng/ml recombinant murine IL-7 (Peprotech, #217-17). For pro-B cells of tet-LIN28B mice, 0.1 µg/ml doxycycline (Merck Life Science, #D3072) was added to induce LIN28B expression. Medium was replaced every other day, and cells were harvested at the indicated time points for flow cytometric analysis. Cell numbers were determined using CountBright™ Absolute Counting Beads (ThermoFisher Scientific, #C36950).

For receptor editing assays, immB cells, including both IgM^low^ and IgM^high^ subsets, were FACS sorted and cultured in B cell medium with Wortmannin (Merck Life Science, #W3144) or F(ab’)2 Anti-Mouse IgM, µ chain specific Functional Grade Purified (eBioscience, #16-5092-85) at the indicated concentrations. To limit cell death associated with prolonged *ex vivo* culture, cells were harvested after 16 hours and used for flow cytometric readout or stored in DNA/RNA shield for subsequent RNA extraction. Cell numbers were determined using CountBright™ Absolute Counting Beads.

### EdU labelling and detection

To trace proliferating cells, mice received a single intraperitoneal injection of 5 mg 5-Ethynyl-2′-deoxyuridine (EdU, Abcam, #ab146186), as previously described (*37*), and were euthanized for analysis 48 hours later. Following BM isolation, cells were stained for surface markers, and EdU incorporation and intracellular markers were detected using the Click-iT™ Plus EdU Alexa Fluor™ 647 Flow Cytometry Assay Kit (ThermoFisher Scientific, #C10634) according to the manufacturer’s instructions.

### Statistical analyses

Statistical analyses and visualizations were performed in Graphpad Prism 10. Statistical tests are specified in the corresponding figure legends.

## Acknowledgments

We thank Taras Kreslavskiy (Karolinska institute, Sweden), Stijn Vanhee (Ghent University), Madelene Dahlgren (Lund University) and Jack Polmear (Lund University) for critical feedback on the manuscript. We thank Taras Kreslavskiy (Karolinska institute, Sweden) for providing B1-8hi mice and Lill Mårtensson-Bopp (University of Gothenburg, Sweden) for providing SLC–/– mice. We thank the Animal Facility at Lund University, and the FACS and Bioinformatics Core Facilities at Lund Stem Cell Center for their support with animal husbandry, and data acquisition and analysis. We thank SciLifeLab, and Center for Translational Genomics (CTG) Lund University for providing sequencing service.

## Funding

Swedish Childhood Cancer Fund (EB)

European Research Council, 715313 and 101125425 (JY)

Knut and Alice Wallenberg Foundation, 2024.0097 (JY)

Swedish Research Council, 2022-00617 (JY)

Swedish Cancer Society, 23-2873 (JY)

Svenska Sällskapet för Medicinsk Forskning, Consolidator grant (JY)

Swedish Government Initiative for Strategic Research Areas – Stem Therapy (FACS and Bioinformatics Core Facility)

SciLifeLab & Wallenberg Data Driven Life Science Program, KAW 2020.0239 (CRC)

## Author contributions

Conceptualization: EB, JY

Experimental work: EB, GM, HÅ, NS, CV

Bioinformatic analyses: EB, SS

Support with bioinformatic analyses: SL, SS, CRC, JEF

Funding acquisition: EB, JY

Writing – original draft: EB, JY

Writing – review & editing: all authors

## SUPPLEMENTARY FIGURES

**Supplementary Figure 1.**
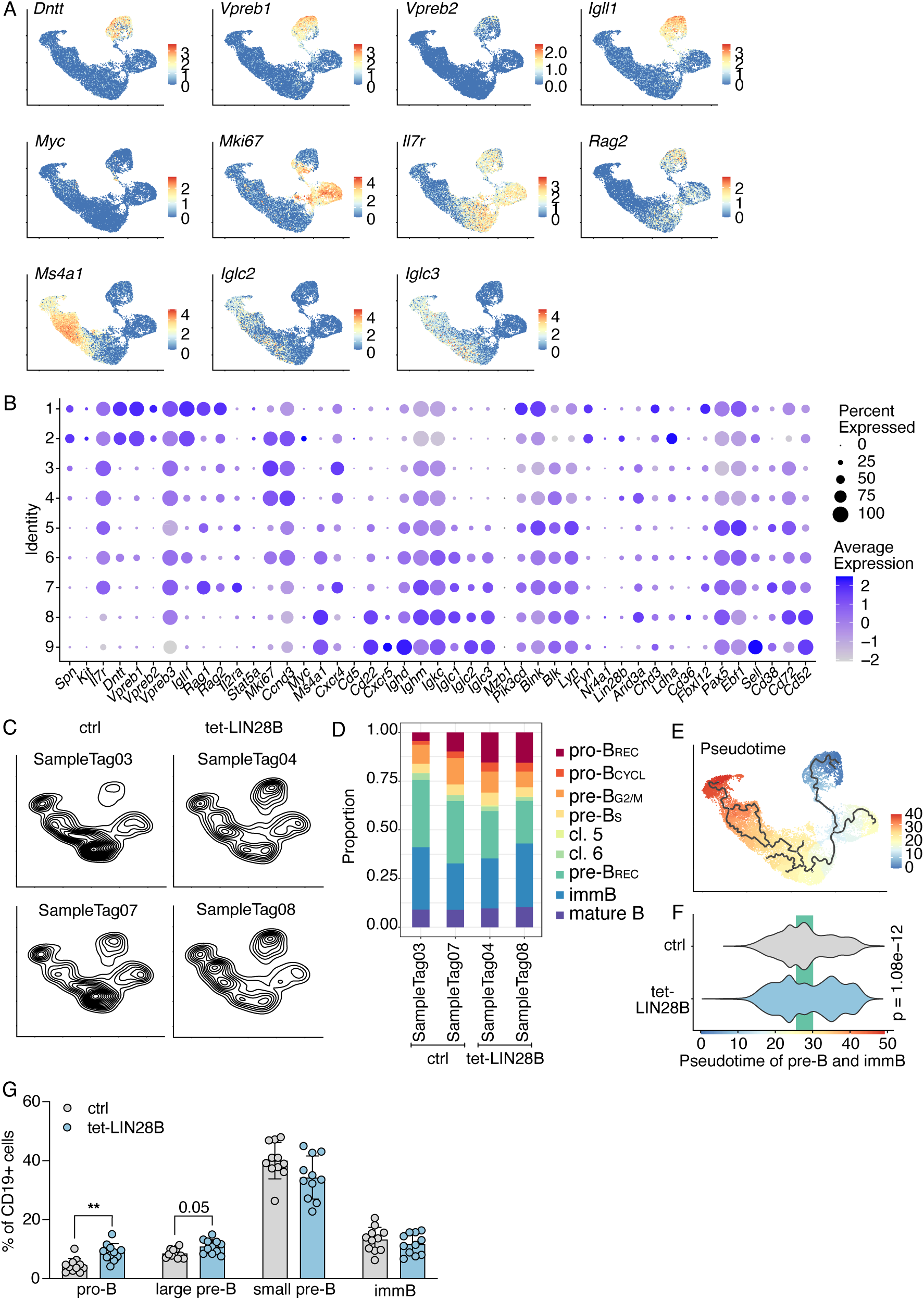
scRNAseq and flow cytometry analysis of B cell progenitors in adult ctrl and tet-LIN28B mice. A. Expression of selected B cell development genes on UMAP representation, according to scRNAseq data. B. Dot plot of the expression of selected B cell development genes in identified Seurat clusters, according to scRNAseq data. Clusters were defined as follows: pro-B cells (*Vpreb1*⁺, *Vpreb2*⁺, *Igllc5*⁺, *Dntt*⁺) were separated in pro-B_REC_ (cluster 1, *Rag1*⁺) and pro-B_CYCL_ (cluster 2, *Mki67*⁺). Pre-B cells (*Vpreb1*^−^, *Vpreb2*^−^, *Igllc5*^−^, *Dntt*^−^, *Il7r*⁺) were divided in pre-B_G2/M_ and pre-B_S_ (*Mki67*⁺, clusters 3 and 4 respectively, phases defined by “CellCycleScoring” in Seurat), pre-B_REC_ (cluster 7, *Rag1*⁺) and two smaller clusters (clusters 5 and 6). ImmB (cluster 8) and recirculating B (cluster 9) cell clusters expressed *Ms4a1*. C. Geometric density of single samples included in scRNAseq analysis. D. Proportion of Seurat clusters in the different samples. E. Monocle3 pseudotime. F. Violin plot of pseudotime in ctrl and tet-LIN28B cells, including stages from cycling pre-B (pre-B_G2/M_ and pre-B_S_) to immB. G. Percentage out of CD19⁺ cells of BCPs in ctrl and tet-LIN28B mice. Pro-B: CD19⁺B220⁺CD93⁺IgM^−^CD117⁺, large pre-B: CD19⁺B220⁺CD93⁺IgM^−^CD117^−^FSC_high_); small pre-B: CD19⁺B220⁺CD93⁺IgM^−^CD117^−^FSClow; immB: CD19⁺B220⁺CD93⁺IgM⁺. Mean ±SD. Two-way ANOVA test, with Tukey correction for multiple comparisons, significant comparisons shown, ** adj.p < 0.01.

**Supplementary Figure 2.**
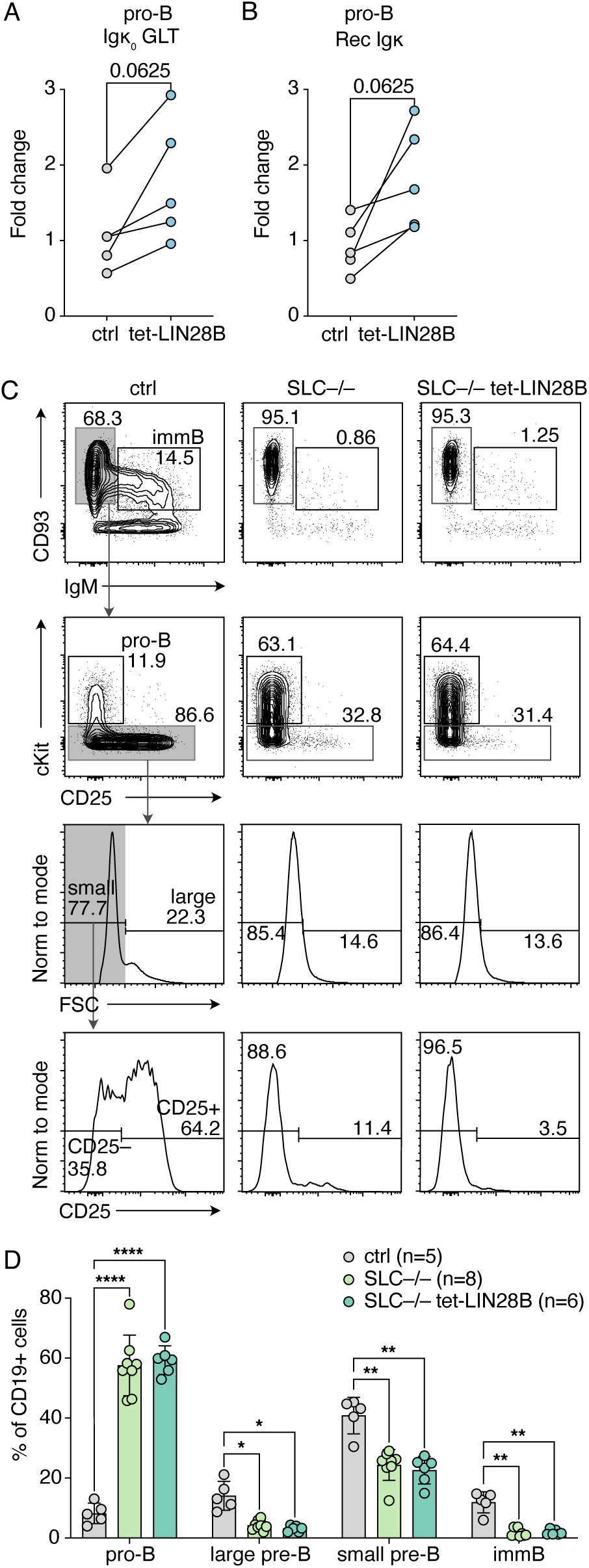
Pre-BCR checkpoint bypass is not a major mechanism of LIN28B action. A. Relative expression of germline transcripts (GLT) of the immunoglobulin κ chain (*Igκ*) in sorted pro-B cells isolated from ctrl (n=5) and tet-LIN28B (n=5) mice. Wilcoxon matched-pairs signed rank test. B. Relative expression of recombined (Rec) *Igκ* in sorted pro-B cells isolated from ctrl (n=5) and tet-LIN28B (n=5) mice. Wilcoxon matched-pairs signed rank test. C. Representative FACS plots of CD19⁺B220⁺ bone marrow cells in ctrl, surrogate light chain deficient (SLC–/–) and SLC–/– tet-LIN28B mice. D. Percentage of B cell progenitors (BCP) in C out of CD19⁺ cells. Mean ±SD. Two-way ANOVA test, with Tukey correction for multiple comparisons, only significant comparisons shown, * adjusted p-value (adj.p) < 0.05, ** adj.p < 0.01, **** adj.p < 0.0001.

**Supplementary Figure 3.**
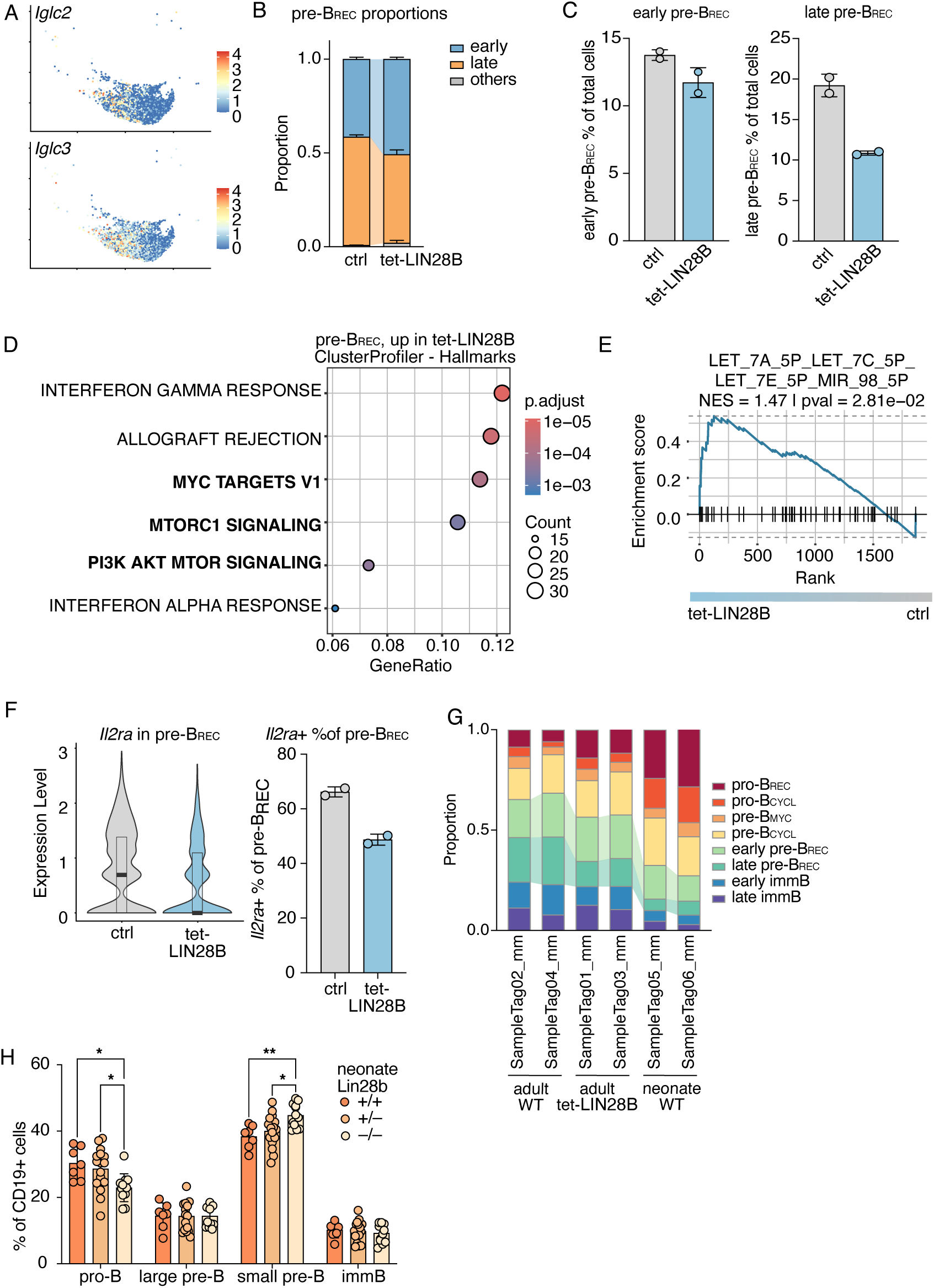
LIN28B-mediated changes on pre-B_REC_ proportions and gene expression. A. *Iglc2* and *Iglc3* expression in pre-B_REC_ cells from scRNAseq experiment. B. Proportion of pre-B_REC_ Seurat clusters in the different genotypes. Mean ±SD. C. Percentage of early and late pre-B_REC_ clusters out of total good-quality cells in the different genotypes. Mean ±SD. D. Results of enrichment for Hallmark Genesets of the MSigDB(*69*) assessed with ClusterProfiler in genes upregulated in tet-LIN28B vs ctrl pre-B_REC_ cells. E. Enrichment of Let-7 target geneset “LET_7A_5P_LET_7C_5P_LET_7E_5P_MIR_98_5P” of the MSigDB(*69*) in tet-LIN28B vs ctrl pre-B_REC_ cells. F. Overlapping box plot (Median ±quartiles) and violin plot of *Il2ra* expression in ctrl and tet-LIN28B pre-B_REC_ cell cluster from scRNAseq experiment, and percentage of immB cells with *Il2ra* expression >0. Mean ±SD. G. Proportion of Seurat clusters for each sample in ctrl adult (n=2), tet-LIN28B adult (n=2), and wildtype (WT) neonate (n=2) mice expressed as proportion out of total cells detected after quality filtering in targeted scRNAseq experiment. H. Percentage out of CD19⁺ cells of BCPs in Lin28b +/+ (n=7), +/– (n=16) and –/– (n=12) neonate mice. Mean ±SD. Two-way ANOVA test, with Tukey correction for multiple comparisons, significant comparisons shown, * adj.p < 0.05, ** adj.p < 0.01.

**Supplementary Figure 4.**
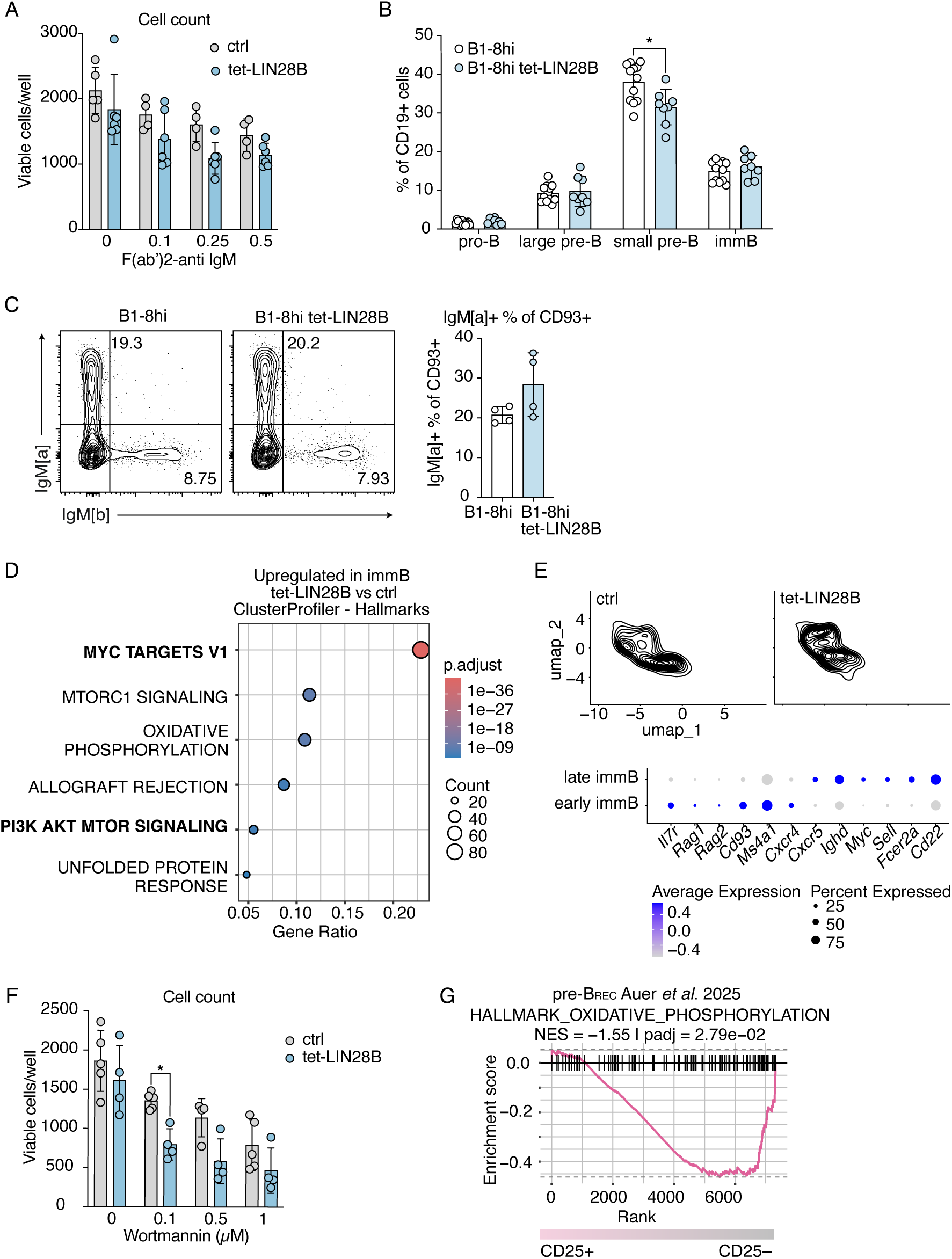
LIN28B represses programs associated with receptor editing. A. Absolute count of viable cells per well in overnight culture of immB cells from ctrl (n=4) or tet-LIN28B (n=6) mice treated with the indicated concentrations of F(ab′)2-anti IgM. Mean ±SD. Two-way ANOVA test, with Šídák correction. B. Percentage out of CD19⁺ cells of B cell progenitors in B1-8hi (n=12) and B1-8hi tet-LIN28B mice (n=8). Two-way ANOVA test, with Šídák correction for multiple comparisons. C. Representative FACS plot and percentage of IgM[a]⁺ cells out of CD19⁺CD93⁺ cells in the BM of B1-8hi mice with and without LIN28B (n=4 per group). Mean ±SD. Mann-Whitney test. D. Enrichment for Hallmark Genesets of the MSigDB(*69*) assessed with ClusterProfiler in genes upregulated in tet-LIN28B vs ctrl immB cells. E. Geometric density of ctrl and tet-LIN28B immB cells from scRNAseq experiment and Dot plot of the expression of selected B cell development genes in early and late immB cells, according to scRNAseq data. F. Absolute count of viable cells per well in overnight culture of immB cells from ctrl (n=5) or tet-LIN28B (n=4) mice treated with the indicated concentrations of Wortmannin. Mean ±SD. Two-way ANOVA test, with Šídák correction. G. Enrichment for “HALLMARK_OXIDATIVE_PHOSPHORYLATION” from the MSigDB(*69*) in CD25⁺ vs CD25^−^ pre-B_REC_ identified reanalyzing a published dataset (*45*) Only significant comparisons shown, * adj.p < 0.05.

## SUPPLEMENTARY TABLES

**Supplementary Table 1. Differentially expressed genes in tet-LIN28B vs ctrl pre-B_REC_, immB, early immB, late immB.** avg_log2FC: log fold-change of the average expression between the two groups; pct: percentage of cells where the gene is detected in each group; p_val_adj: adjusted p-value, based on bonferroni correction using all genes in the dataset.

**Supplementary Table 2. Gene panel for targeted scRNAseq** (***47***).

**Supplementary Table 3. Differentially expressed genes in CD25⁺ vs CD25^−^ pre-B_REC_ in Auer *et al.***(***45***).

**Supplementary Table 4. Genes negatively correlated with *Il2ra* in ctrl samples in WTA scRNAseq data. Spearman correlation < -0.1.**

**Supplementary Table 5.**
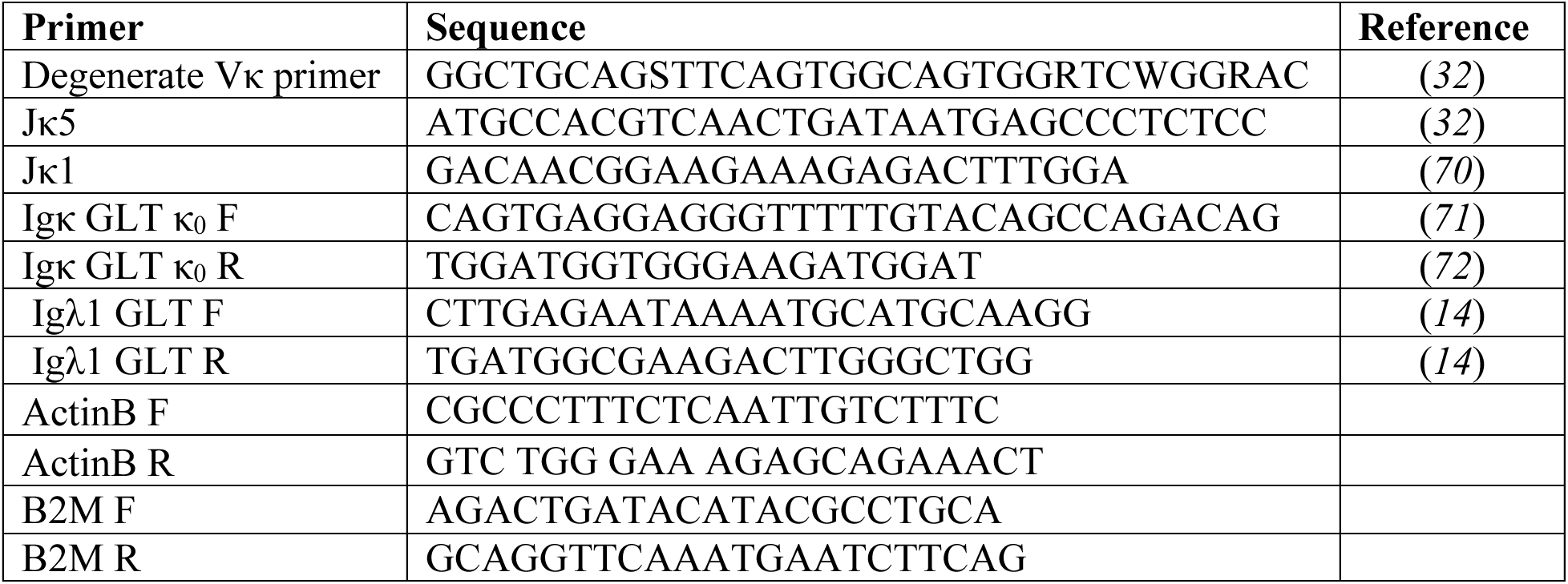
Primer list. GLT: germline transcript; F: forward; R: reverse.

